# Mechanism of PEX5-mediated protein import into peroxisomes

**DOI:** 10.1101/2022.05.31.494222

**Authors:** Michael L. Skowyra, Tom A. Rapoport

## Abstract

Peroxisomes are ubiquitous organelles, whose dysfunction causes fatal human diseases. Most peroxisomal enzymes are imported from the cytosol by the receptor PEX5, which interacts with a docking complex in the peroxisomal membrane, and then returns to the cytosol after monoubiquitination by a membrane-embedded ubiquitin ligase. The mechanism by which PEX5 shuttles between cytosol and peroxisomes, and releases cargo inside the lumen, is unclear. Here, we use *Xenopus* egg extract to demonstrate that PEX5 accompanies cargo completely into the lumen, utilizing WxxxF/Y motifs near its N-terminus that bind a lumenal domain of the docking complex. PEX5 recycling is initiated by an amphipathic helix that binds to the lumenal side of the ubiquitin ligase. The N-terminus then emerges in the cytosol for monoubiquitination. Finally, PEX5 is extracted from the lumen, resulting in unfolding of the receptor and cargo release. Our results reveal the unique mechanism by which PEX5 ferries proteins into peroxisomes.

## Introduction

Peroxisomes are membrane-bounded organelles that exist in nearly all eukaryotic cells (Jansen et al., 2021), and provide a number of vital metabolic functions including the oxidation of fatty acids and other biomolecules (Islinger et al., 2010; Kao et al., 2018). Because many of these reactions generate hydrogen peroxide and other reactive oxygen species, peroxisomes additionally house catalase and other enzymes involved in redox homeostasis (Lismont et al., 2019). In humans and other animals, they are also required for the production of bile salts and of lipids contained in myelin (Ferdinandusse et al., 2009; Kassmann, 2014). The importance of peroxisomes to human health is revealed by various disorders, notably the Zellweger spectrum, that are caused by defects in peroxisome biogenesis and are often fatal (Fujiki, 2016). Peroxisome biogenesis disorders frequently arise from inherited mutations in proteins called peroxins (PEX) that are responsible for importing enzymes into the peroxisomal lumen, or matrix (Steinberg et al., 2006).

Matrix proteins are synthesized in the cytosol and delivered post-translationally to peroxisomes, which in humans and animals is mediated exclusively by the soluble receptor PEX5 (for review, see Farré et al., 2019; Francisco et al., 2017; Fujiki et al., 2014; Walter and Erdmann, 2019). PEX5 consists of a long unstructured N-terminal region of poorly defined function, followed by a globular tetratricopeptide repeat (TPR) domain (Barros-Barbosa et al., 2018). The TPR domain directly binds the PTS1 peroxisomal targeting signal, which occurs in most matrix proteins and is composed of a C-terminal Ser-Lys-Leu (SKL) tripeptide or variants of it (Gatto et al., 2000). A few matrix proteins contain an alternative N-terminal signal called PTS2, whose recognition requires the adapter PEX7 (Pan et al., 2013). In higher organisms, PEX7 associates with PEX5 through a motif in the receptor’s unstructured N-terminal region (Dodt et al., 2001; Nito et al., 2002). A shorter version of PEX5 lacking this motif is produced by alternative splicing and can only bind PTS1 cargo (Braverman et al., 1998; Otera et al., 1998).

Cargo-loaded PEX5 is recruited to peroxisomes by a docking complex, which includes the membrane proteins PEX13 and PEX14 (for review, see Azevedo and Schliebs, 2006; Kalel and Erdmann, 2018; Kao et al., 2018; Williams and Distel, 2006). *In vitro*, PEX5 binds the docking components using conserved aromatic residues in WxxxF/Y pentapeptide motifs (where “x” denotes any amino acid) located within its N-terminal unstructured region (Nito et al., 2002; Otera et al., 2002; Saidowsky et al., 2001; Urquhart et al., 2000). Whether these motifs mediate binding of PEX5 to the cytosolic surface of peroxisomes or participate in a later step is unclear, given that there are conflicting reports on the cytosolic or lumenal localization of the cognate binding domains in the docking complex (reviewed in Barros-Barbosa et al., 2018). Docking is followed by translocation of cargo into the peroxisomal lumen by a poorly understood process, which allows even folded or oligomeric proteins to traverse the membrane (Dias et al., 2016; Léon et al., 2006). In this respect, the mechanism of translocation into peroxisomes must differ dramatically from that into the endoplasmic reticulum (ER) or mitochondria, which can only import proteins in an unfolded conformation (Dudek et al., 2013; Rapoport et al., 2017).

To complete the import cycle, PEX5 must return to the cytosol. This recycling step requires ubiquitination of PEX5 by a membrane-embedded, RING-type ubiquitin ligase complex composed of PEX2, PEX10, and PEX12 (for review, see Francisco et al., 2014; Kao et al., 2018; Wang and Subramani, 2017). Curiously, PEX5 appears to be monoubiquitinated at a conserved cysteine near the N-terminus (Carvalho et al., 2007; Platta et al., 2007; Williams et al., 2007), an unusual residue to be modified. Monoubiquitinated PEX5 is thought to be extracted from peroxisomes by a hexameric AAA ATPase formed from alternating copies of PEX1 and PEX6 (Blok et al., 2015; Miyata and Fujiki, 2005; Platta et al., 2005). The ubiquitin moiety is then likely removed in the cytosol by de-ubiquitinating enzymes (Debelyy et al., 2011; Grou et al., 2012), allowing PEX5 to initiate a new round of cargo translocation.

The role of PEX5 in moving cargo across the peroxisomal membrane is a longstanding question. According to one model, PEX5 would form an integral component of a translocation channel (Erdmann and Schliebs, 2005), possibly through association with PEX14 (Meinecke et al., 2010). This model is supported by observations of PEX5 adopting a transmembrane topology in the peroxisomal membrane, with the N-terminus facing the cytosol (Azevedo and Schliebs, 2006). Although this topology would explain how the N-terminal cysteine is modified by ubiquitination machinery localized in the cytosol, the absence of hydrophobic elements in PEX5 befogs how the receptor would integrate into the membrane. According to an alternative model, PEX5 would shuttle cargo into the peroxisomal lumen (Dammai and Subramani, 2001; Kunau, 2001). This model is supported by observations of the receptor’s N-terminus penetrating into the lumen during import (Dammai and Subramani, 2001), and by experiments suggesting at least partial lumenal entry of receptor-bound PEX7 (Nair et al., 2004). However, it is unclear whether the entire receptor moves across the membrane. Lumenal entry of PEX5 also raises the question of how the receptor, in particular the folded TPR domain, would return to the cytosol. Importantly, neither model can easily explain how cargo would be released in the lumen. Given the conflicting results in the literature, the mechanism by which PEX5 mediates protein import into peroxisomes remains unresolved.

Here, using a cell-free system based on *Xenopus* egg extract, we demonstrate that PEX5 accompanies cargo all the way into peroxisomes and then returns to the cytosol. We identify the features in PEX5 required for admission into the lumen and for subsequent egress, and show that PEX5 enters the organelle in a folded state but is unfolded during retro-translocation. These results reveal that PEX5 functions as a shuttle, and suggest a mechanism for cargo release inside the peroxisomal matrix.

## Results

### An *in vitro* assay that recapitulates PEX5 cycling during peroxisomal protein import

We previously described a cell-free system based on *Xenopus* egg extract, which reproduces matrix protein import into peroxisomes (Romano et al., 2019). Extract corresponds to the cytoplasm of eggs from the African clawed frog, *Xenopus leavis*, and is generated by a gentle procedure (Fig. 1A) that preserves organellar integrity and maintains cellular components at near physiological concentrations. Protein import can be visualized in the extract using fluorescent proteins fused to a canonical PTS1 signal (Fig. 1A). The accumulation of the fusion protein in bright puncta can be quantitatively followed over time by fluorescence microscopy (panel 1). This system allows components to be acutely manipulated and replaced with mutant versions, minimizing indirect effects that may occur in live cells.

**Figure 1.**
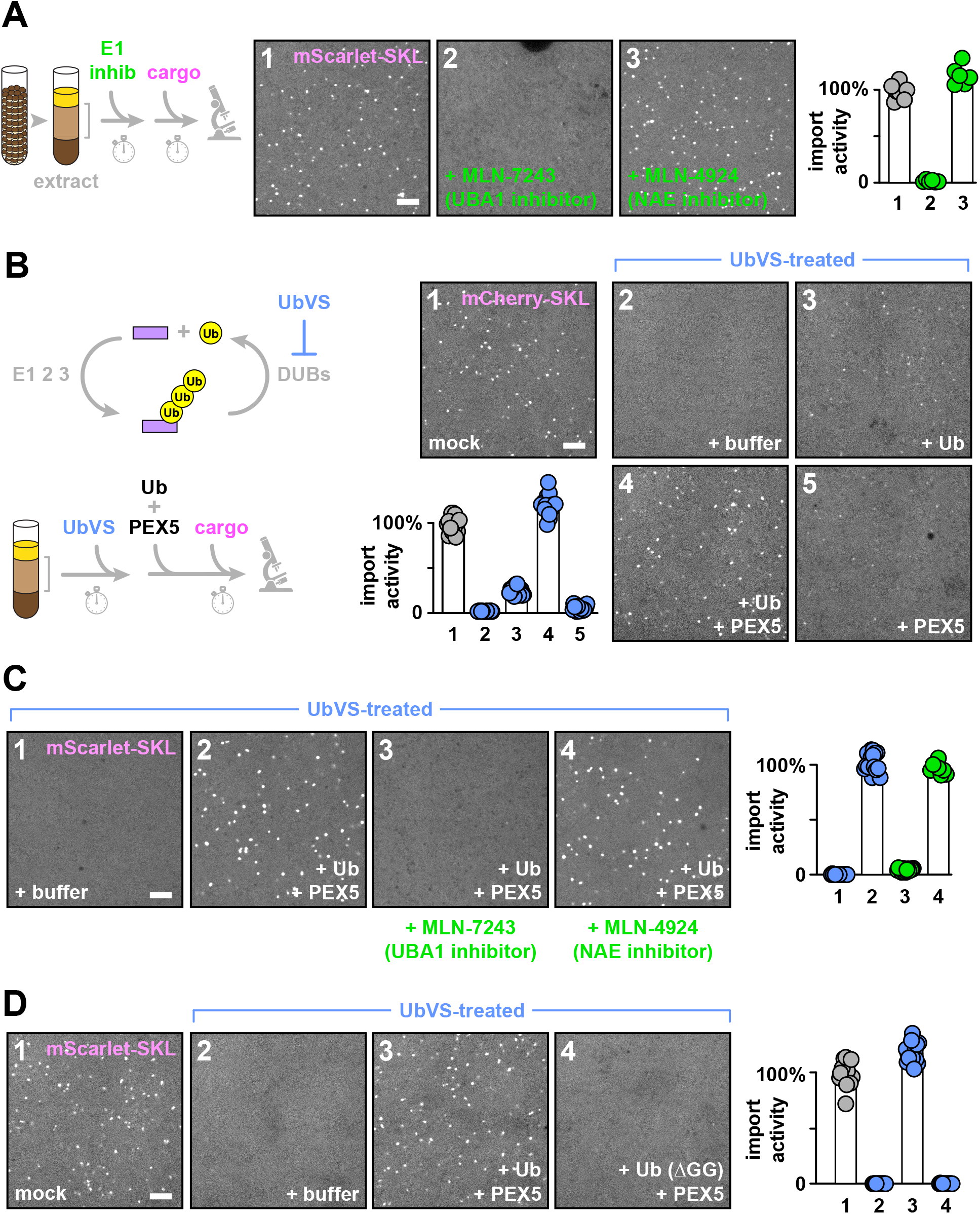
PEX5 recycling recapitulated in *Xenopus* egg extract. (A) *Xenopus* eggs were lysed by centrifugation to generate an extract (the indicated layer between the lipid on top and cellular debris on bottom). The extract was treated with DMSO or the indicated E1 enzyme inhibitors dissolved in DMSO (UBA1, ubiquitin-activating enzyme; NAE, NEDD8-activating enzyme). Import activity was then assessed by incubating with the fluorescent cargo mScarlet-SKL and imaging on a spinning disk confocal microscope; repeated rounds of import into peroxisomes result in the appearance of bright puncta. The number of puncta in an imaged field was quantified relative to that in the untreated reaction (*n* = 9 fields per reaction; bars specify the median). (B) Upper scheme illustrates the strategy to deplete free ubiquitin from extract by inhibiting deubiquitinating enzymes (DUBs) with ubiquitin vinyl sulfone (UbVS); lower scheme shows the approach for restoring peroxisome import activity. On the right, extract was treated with buffer (mock) or UbVS, then supplemented either with buffer or wild type ubiquitin, wild type PEX5, or both components together. Import activity was then assessed as above (*n* = 18 fields per reaction). (C) As in (B), except that reactions were supplemented with the indicated components in the absence or presence of the E1 inhibitors. Import activity was assessed as above (*n* = 18 fields per reaction). (D) As in (C). Ubiquitin mutant (ΔGG) lacks both C-terminal glycines. Import activity was assessed as above (*n* = 18 fields per reaction). Scale bars equal 5 µm. See also supplemental figure S1.

Using this *in vitro* system, we tested whether PEX5 cycles between the cytosol and peroxisomes to mediate multiple rounds of matrix protein import. Because PEX5 must be ubiquitinated every time it returns to the cytosol, sustained import should depend on ubiquitination. Indeed, import was greatly attenuated in extract treated with MLN-7243 (Hyer et al., 2018), an inhibitor of the primary ubiquitin-activating (E1) enzyme UBA1 (Fig. 1A, panel 2). In contrast, no effect on import was observed with an inhibitor that targets a related E1, the NEDD8-activating enzyme (NAE) (Brownell et al., 2010) (Fig. 1A, panel 3). As a complementary approach, we depleted free ubiquitin from extract by inhibiting de-ubiquitinating enzymes (DUBs) with ubiquitin vinyl sulfone (UbVS) (Fig. 1B); this treatment causes sequestration of free ubiquitin in conjugates so that it becomes unavailable for further reactions (Dimova et al., 2012). As expected, import was considerably reduced (panel 2 versus 1). Since a single round of import does not require ubiquitination, these data indicate that multiple rounds of import occur in our system.

Next, we attempted to restore import activity in UbVS-treated extract. Addition of purified ubiquitin could partially restore import (Fig. 1B, panel 3), but complete rescue could not be achieved even with a large excess over the endogenous levels (**fig. S1A**). Such high concentrations did not compromise import in untreated extract (**fig. S1B**), excluding the presence of an inhibitory contaminant. Instead, the partial rescue raised the possibility that PEX5 had become limiting, as, in the absence of DUB activity, it might have been converted into a ubiquitinated form unable to mediate import. We therefore supplemented UbVS-treated extract with ubiquitin together with an excess of purified recombinant PEX5 over the endogenous receptor (the purity of the PEX5 proteins is shown in **fig. S1D**). Import was fully restored (Fig. 1B, panel 4), whereas PEX5 alone had little effect (Fig. 1B, panel 5). Similar results were obtained by depleting ubiquitin with other DUB inhibitors (**fig. S1C**). Confirming the need for ubiquitin conjugation, inhibition of UBA1 precluded the rescue of import activity by ubiquitin and PEX5, whereas inhibition of NAE had no effect (Fig. 1C). Furthermore, a ubiquitin mutant lacking the terminal two amino acids (ΔGG) could not replace wild type ubiquitin in this assay (Fig. 1D). Taken together, these results confirm that PEX5 ubiquitination is required for sustained peroxisomal protein import.

Next, we tested whether sustained import depends on the reported monoubiquitination of PEX5 at a conserved cysteine (Cys11 in vertebrates). Methylated ubiquitin or a variant lacking all lysines, neither of which can form ubiquitin chains, restored as much import as wild type ubiquitin when added together with PEX5 (Fig. 2A), demonstrating that polyubiquitination is dispensable. Furthermore, import was not restored when a PEX5 mutant was used that lacked the conserved cysteine (C11A), whereas a lysine at this position (C11K) retained full activity (Fig. 2B), as reported previously (Grou et al., 2009; Romano et al., 2019). To test whether the conserved cysteine is the only ubiquitination site required for import, we depleted endogenous PEX5 with antibodies and replaced it with different mutants (Fig. 2C). A mutant lacking all lysines (ΔK) was still active (panel 4), as was a mutant that contained only a single lysine at position 11 (ΔK + C11K) (panel 6). Blocking ubiquitination of this lysine by methylation abolished import (panel 7), whereas PEX5 (ΔK) containing the native cysteine remained active after methylation (panel 5). These results indicate that monoubiquitination is sufficient for the import function of PEX5, and establish that the conserved cysteine is the only ubiquitination site required. The dependence on ubiquitination confirms that PEX5 cycles between the cytosol and peroxisomes to mediate multiple rounds of protein import in our cell-free system.

**Figure 2.**
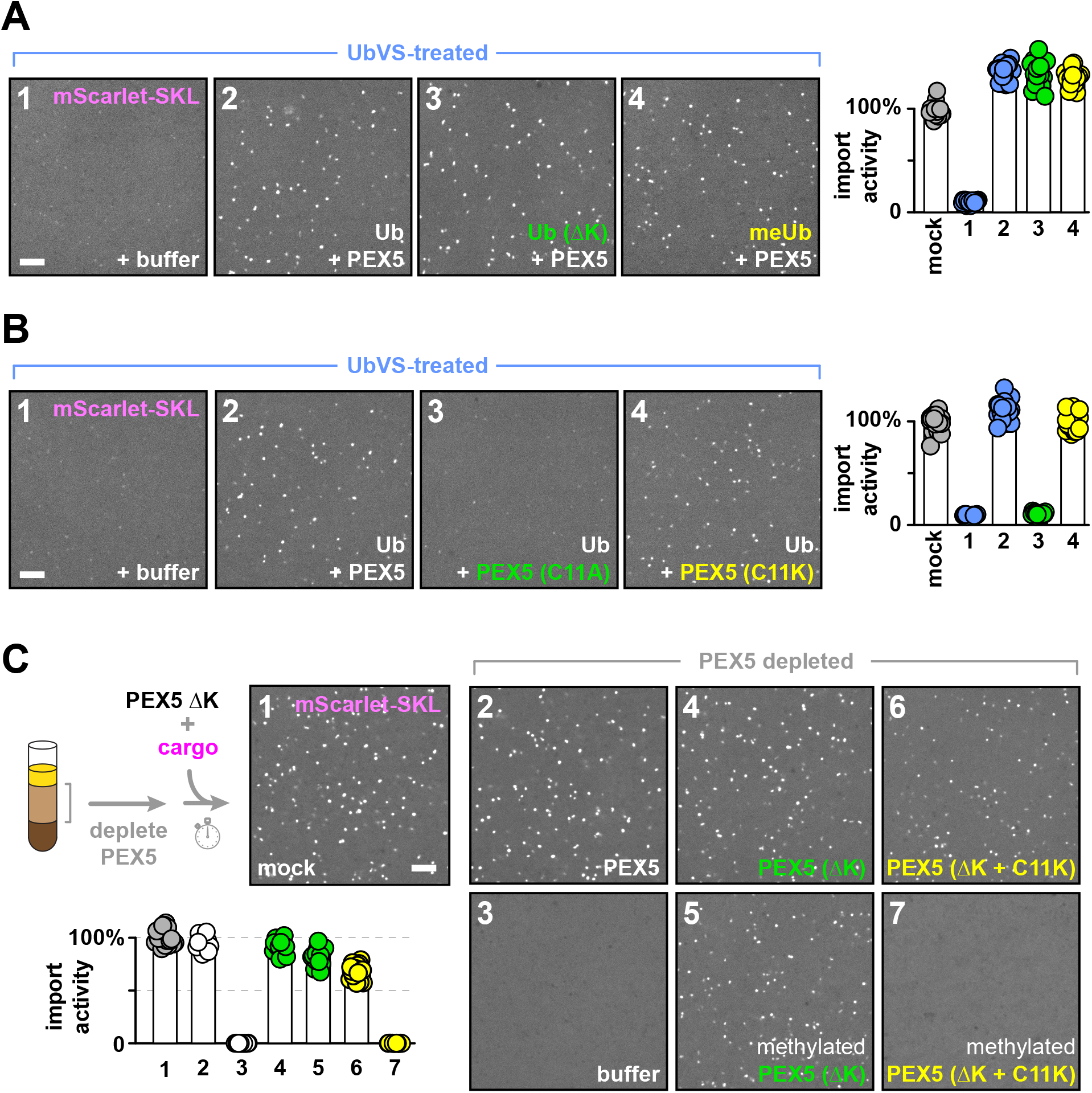
Sustained peroxisomal import requires monoubiquitination and the conserved cysteine in PEX5. (A) *Xenopus* egg extract was treated with UbVS, then supplemented either with buffer or PEX5 together with wild type ubiquitin (Ub), ubiquitin lacking all lysines (Ub ΔK), or methylated ubiquitin (meUb). Import activity was assessed by incubating with mScarlet-SKL and imaging the formation of bright puncta on a spinning disk confocal microscope. The number of puncta in an imaged field was quantified relative to that in the mock reaction (*n* = 18 fields per reaction; bars specify the median). (B) As in (A), except that reactions were supplemented with ubiquitin together with wild type PEX5 or PEX5 mutants in which Cys11 was converted to Ala (C11A) or Lys (C11K). Import activity was assessed as above (*n* = 18 fields per reaction). (C) Extract was depleted of endogenous PEX5 using beads conjugated to PEX5 antibodies, then supplemented either with buffer or wild type PEX5, a PEX5 mutant lacking all lysines (ΔK), or PEX5 ΔK in which Cys11 was mutated to Lys (ΔK + C11K). Reactions 5 and 7 were performed with methylated versions of the indicated mutants. Import activity was assessed as above (*n* = 18 fields per reaction). All scale bars equal 5 µm. See also supplemental figure S2.

Interestingly, peroxisomal import was unaffected by replacing endogenous ubiquitin in extract with mutants (F4A, I44A, D58A, or V70A) known to block other ubiquitin-dependent processes (**fig. S2A**). In contrast, ablating residues in ubiquitin close to the C-terminal tail (L8 or L73) inhibited import (**fig. S2A**) without compromising the conjugation of the corresponding mutants to the E2 enzyme implicated in PEX5 recycling (**fig. S2B**). These results suggest that the ubiquitin molecule in monoubiquitinated PEX5 interacts with the recycling machinery, likely the uniquitin ligase complex, using a surface proximal to the C-terminal tail. A similar surface is important for ubiquitination by other RING-type ligases (Plechanovová et al., 2012).

### Signals targeting PEX5 to peroxisomes

Next, we sought to identify the signals in PEX5 that direct the receptor to peroxisomes. Previous work suggested that these signals are located within the N-terminal unstructured region (Dodt et al., 2001). In humans and other animals, this region contains multiple WxxxF/Y pentapeptide motifs (designated W1-W7 in Fig. 3A) along with an analogous LxxxF motif (designated W0). The importance of these pentapeptide motifs has been recognized before (Neuhaus et al., 2014; Otera et al., 2002), although their individual contributions and exact function remain unclear. The N-terminal region additionally includes four predicted amphipathic helices (designated AH1-AH4 in Fig. 3A) of unknown significance (Gaussmann et al., 2021). AH1 resides downstream of Cys11 and has a hydrophobic surface conserved across eukaryotes (Fig. 3B). An uncharacterized asparagine (Asn15) precedes AH1 in most PEX5 homologs (**fig. S3A**). AH2 and the other amphipathic helices have hydrophobic surfaces that are conserved in animals (Fig. 3B **and fig. S4A**), and helices with similar features may exist in other organisms (**fig. S4B**).

**Figure 3.**
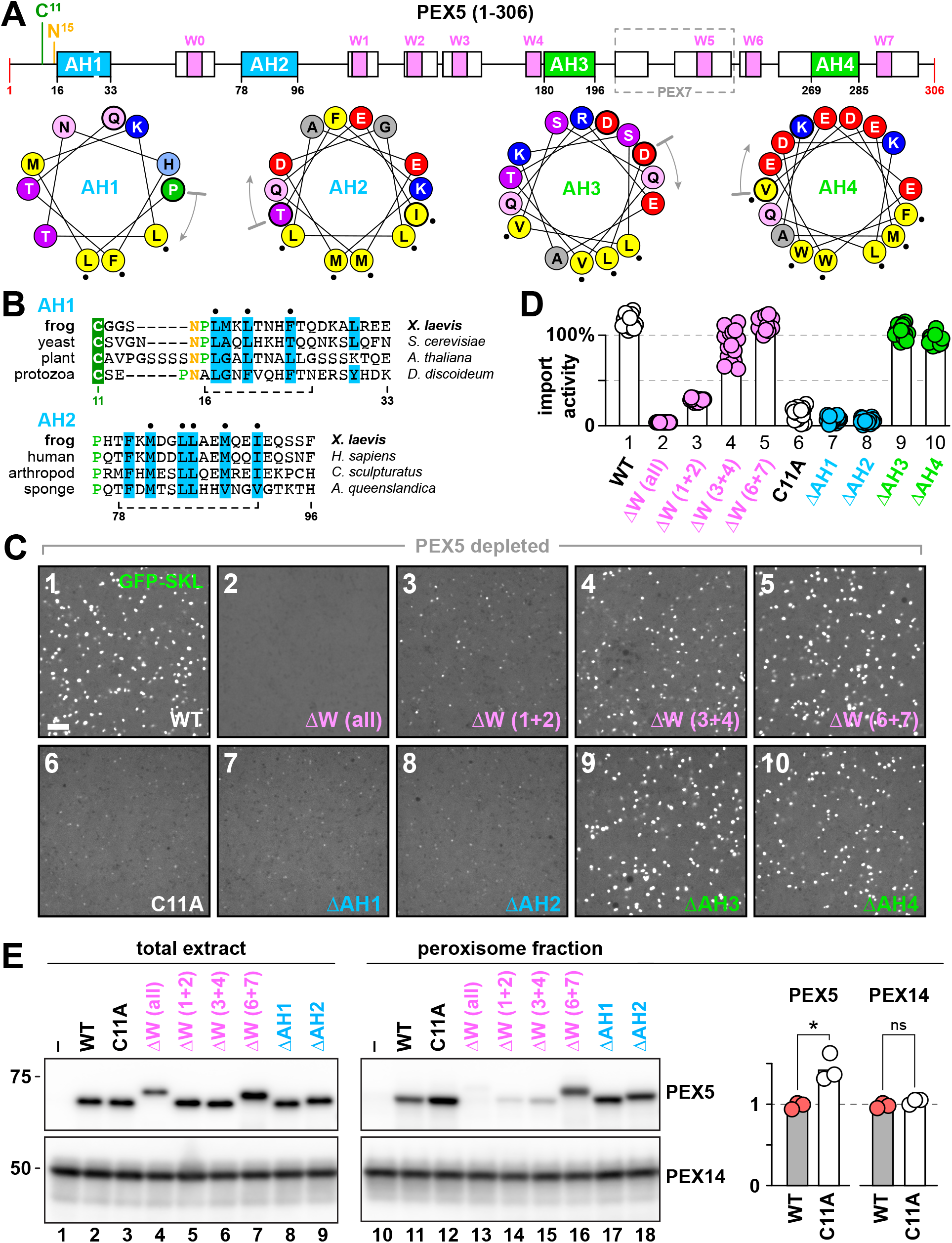
Signals targeting PEX5 to peroxisomes. (A) Diagram on top shows the locations of predicted α-helices (boxes) and key residues in the N-terminal unstructured region of *X. laevis* PEX5 (AH1-AH4, amphipathic helices; W0-W7, WxxxF/Y motifs). The PEX7-binding region is enclosed by a gray dashed line. Helical wheels beneath the diagram illustrate the distribution of hydrophobic amino acids in each amphipathic helix. Gray arrows specify the N-terminus of each helix; black dots denote residues that were mutated to alanines. (B) Sequence alignments of AH1 and AH2 in PEX5 homologs from the indicated organisms. Residue numbers refer to *X. laevis* PEX5; dashed line specifies the region shown in the helical wheels (see figure S3A for alignments of AH3 and AH4). (C) *Xenopus* egg extract was depleted of endogenous PEX5 using beads conjugated to the PEX5-binding domain from PEX14, then supplemented with wild type PEX5 (WT) or the indicated mutants (C11A, Cys11 converted to Ala; ΔW, the specified WxxxF/Y motifs mutated to AxxxA; ΔAH1-ΔAH4, mutations in the hydrophobic face of each amphipathic helix as described above). Import activity was assessed by incubating with GFP-SKL and imaging the formation of bright puncta by spinning disk confocal microscopy. The number of puncta in an imaged field was quantified relative to that in the reaction with wild type PEX5 (*n* = 18 fields per reaction; bars specify the median). Scale bar equals 5 µm. (D) Quantification of experiments shown in (C). (E) Extract was incubated with the indicated PEX5 mutants and GFP-SKL, and peroxisomes were then isolated by flotation. Total extract (0.01%) and the peroxisome-containing fraction (2%) were immunoblotted for PEX5 and PEX14. Relative molecular weights (kD) are marked on the left. Amounts of PEX5 and PEX14 in the peroxisome fraction were quantified by densitometry (*n* = 3 independent experiments; *, *p* ≤ 0.01 by Student’s unpaired two-tailed *t*-test). See also supplemental figures S3-S6, and S7.

We first examined the role of the pentapeptide motifs W0-W7. The motifs were mutated to AxxxA, either individually or in combination, and the resulting mutants were added to extract depleted of the endogenous receptor. Depletion was accomplished using beads conjugated to the N-terminal domain of PEX14 (Romano et al., 2019), which binds strongly to the pentapeptide motifs (Saidowsky et al., 2001). To simplify the analysis, the experiments were performed with the short splice variant of PEX5 that lacks the PEX7-binding region and motif W5 (Fig. 3A). Addition of this isoform to PEX5-depleted extract fully restored import of PTS1 cargo (**fig. S5A**), indicating that motif W5 is not needed. Import activity of PEX5 was abolished when all remaining pentapeptide motifs were mutated (Fig. 3C and D), confirming their importance. None was essential when mutated individually (**fig. S5B**), although a slight reduction in activity was observed in the absence of some N-terminal motifs. Ablating two N-terminal motifs together considerably reduced import (Fig. 3C and D), whereas little effect was observed after mutating combinations of more downstream motifs. Consistent with these phenotypes, PEX5 mutants containing individual motifs were all inactive (**fig. S6A**), while mutants bearing combinations of the LxxxF motif (W0) and N-terminal WxxxF/Y motifs could restore some activity (**fig. S6B**). We therefore conclude that import requires the LxxxF motif and WxxxF/Y motifs located near the N-terminus, whereas the more distal motifs W5-W7 are dispensable.

Next, we evaluated the role of the amphipathic helices. For each helix, we converted all conserved amino acids on the hydrophobic face into alanines (Fig. 3A). Import was abolished after mutating either AH1 or AH2, revealing that each of these two helices is vital (Fig. 3C and D). Import was also diminished, but not eliminated, after ablating the conserved Asn preceding AH1 (**fig. S3B**). On the other hand, mutating AH3 or AH4 had no effect (Fig. 3C and D), nor did deletion of a larger segment that encompasses AH4 (**fig. S3C**). Import was also unaffected after combining this deletion with mutations in the AH3 hydrophobic face (**fig. S3C**), excluding possible redundancy of AH3 and AH4. Taken together, this analysis shows that PEX5 relies on the amphipathic helices and pentapeptide motifs in the N-terminal half of its unstructured region to import cargo into peroxisomes.

We used a flotation assay to determine which of these signals is required for recruiting PEX5 to peroxisomes. After centrifuging egg extract in a discontinuous sucrose gradient, peroxisomes containing a pre-imported fluorescent protein accumulated in the top fractions, together with the membrane-associated docking component PEX14 (**fig. S7A**). Cytosol containing unimported material stayed at the bottom. As expected from the role of PEX5 in protein import, a population of the endogenous receptor co-floated with peroxisomes, while the rest remained in the cytosol fractions (**fig. S7A**). Supplementing extract with recombinant PEX5 increased the peroxisome-bound population (Fig. 3E, compare lanes 10 and 11) and accelerated import (**fig. S7B**), indicating that translocation sites are normally not saturated. A PEX5 mutant lacking Cys11 (i.e., C11A) accumulated to a significantly greater extent on peroxisomes (Fig. 3E, compare lanes 11 and 12, and corresponding quantitation), consistent with this mutant’s inability to return to the cytosol. In contrast, mutating the pentapeptide motifs drastically reduced binding to peroxisomes (Fig. 3E, lanes 13-16), with loss of the N-terminal motifs being more detrimental than loss of downstream ones, consistent with the import activities of these mutants. Thus, the pentapeptide motifs are necessary for recruiting PEX5 to peroxisomes. Notably, amphipathic helices AH1 and AH2 were not required for peroxisomal targeting (Fig. 3E, lanes 17-18); they must therefore play a role at a step downstream from docking.

While PEX5’s pentapeptide motifs are known to bind strongly to the N-terminal domain of PEX14 (Saidowsky et al., 2001), conflicting results exist on the localization of this domain (reviewed in Barros-Barbosa et al., 2018). To clarify this issue, we analyzed the membrane topology of PEX14 in the *Xenopus* system by protease protection (**fig. S7C**). Using an antibody raised against the N-terminal domain, a protease-protected fragment was observed that is consistent with the N-terminus residing in the lumen. This fragment was completely degraded in the presence of detergent. In agreement with the domain’s lumenal orientation, no protease-protected fragments were discernible with an antibody raised against the C-terminus of PEX14. These data therefore suggest that another region of the docking complex mediates the initial interaction between PEX5’s pentapeptide motifs and the cytosolic surface of peroxisomes. Importantly, the strong interaction between the lumenal domain of PEX14 and the pentapeptide motifs also implies that, at some point during the import cycle, PEX5 enters the lumen.

### PEX5 is first imported into peroxisomes and then exported

To determine whether docking is followed by entry of PEX5 into the lumen, we examined the peroxisome-associated population of PEX5 by protease protection. After supplementing extract with wild type PEX5, approximately half of the peroxisome-bound pool was degraded at a low concentration of proteinase K (Fig. 4A), likely reflecting PEX5 molecules docked on the outside of the organelle. The rest of the population was more resistant, but was converted into a slightly shorter form at a high proteinase K concentration (Fig. 4A); all Pex5 molecules were degraded in the presence of detergent. Thus, during cycling between the cytosol and peroxisomes, a short segment of PEX5 becomes exposed to the cytosol while the rest of the protein has moved across the membrane, indicating that the receptor adopts a transmembrane topology. Similar results were obtained with the non-recyclable C11A mutant (Fig. 4A). Consistent with the increased accumulation of this mutant on peroxisomes, the truncated population was also more abundant (Fig. 4A **and fig. S7D**).

**Figure 4.**
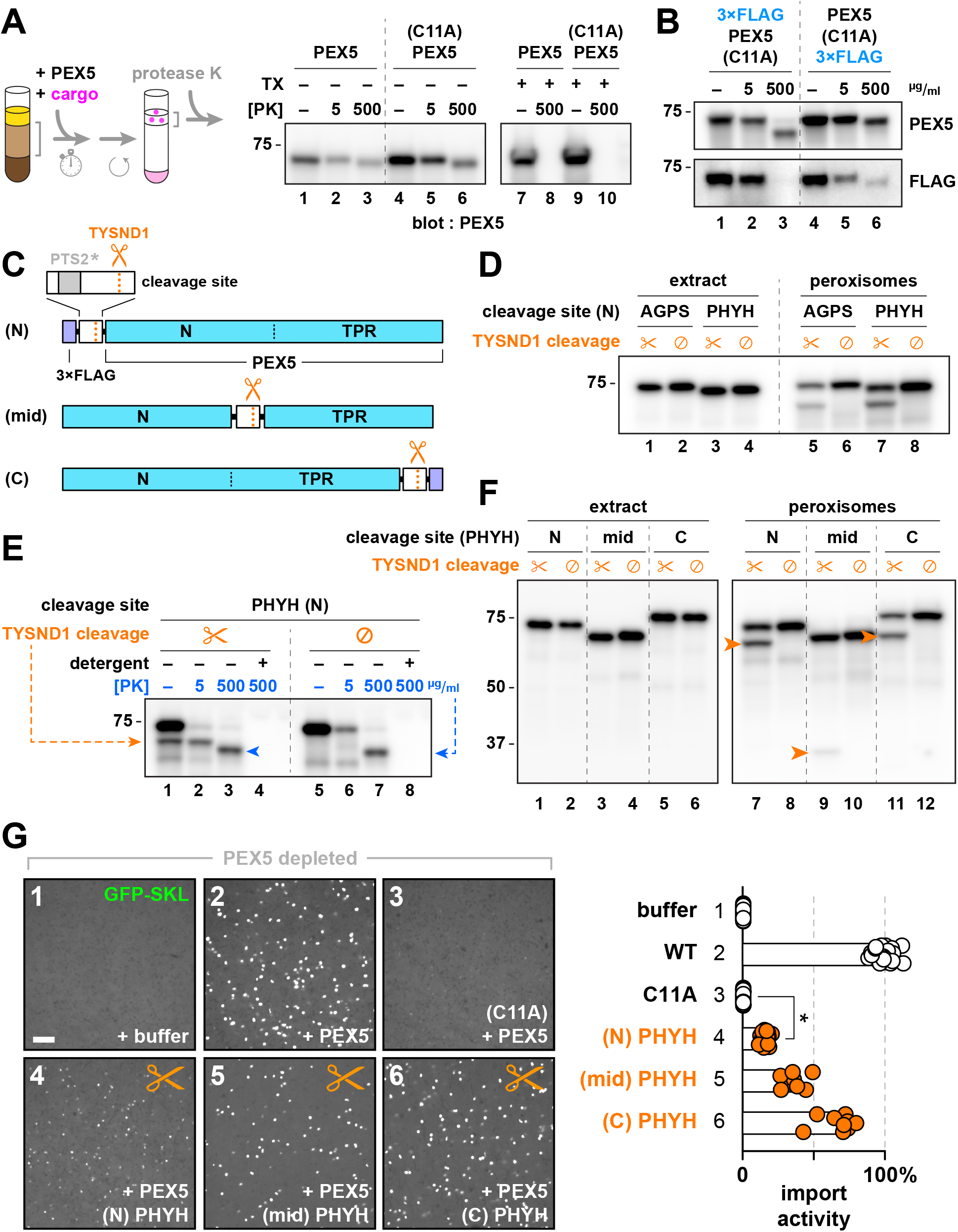
PEX5 export from the peroxisomal lumen. (A) *Xenopus* egg extract was incubated with wild type PEX5 or a mutant lacking the conserved cysteine (C11A) in the presence of GFP-SKL. The peroxisome-associated population was then isolated by flotation and treated with different concentrations of proteinase K, with or without Triton X-100 (TX), and immunoblotted for PEX5. Relative molecular weights (kD) are marked on the left. See also figure S7D. (B) As in (A), except using PEX5 (C11A) mutants fused to a triple FLAG (3×FLAG) tag at either terminus. (C) Scheme depicting the constructs containing cleavable PTS2 signals, which were used to assess whether PEX5 enters the lumen during import. Scissors indicate the site of cleavage by the peroxisomal lumenal protease TYSND1; PTS2* denotes an inactivated PTS2 motif. (D) Extract was incubated with GFP-SKL and a PEX5 (C11A) fusion containing an N-terminal TYSND1 cleavage site from the matrix enzymes AGPS or PHYH (construct (N) in panel C), and peroxisomes were then isolated by flotation. Total extract (0.01%) and the peroxisome-containing fraction (2%) were immunoblotted for PEX5. Scissors denote constructs with an intact TYSND1-cleavage site, whereas (∅) denotes an inactivated site. Relative molecular weights (kD) are marked on the left. (E) As in (D), except using only the construct with the TYSND1-cleavage site from PHYH. Peroxisomes isolated by flotation were treated with different concentrations of proteinase K, with or without Triton X-100 (detergent), and immunoblotted for PEX5. (F) As in (D), except using PEX5 (C11A) containing the TYSND1-cleavage site from PHYH at the indicated positions, as illustrated in (C). (G) Extract was depleted of endogenous PEX5 using beads conjugated to the PEX5-binding domain from PEX14, and supplemented with wild type PEX5, PEX5 (C11A), or wild type PEX5 containing the PHYH TYSND1-cleavage site at the indicated positions (as illustrated in (C)). Import activity was assessed by incubating with GFP-SKL and imaging the formation of bright puncta by spinning disk confocal microscopy. The number of puncta in an imaged field was quantified relative to that in the reaction with wild type PEX5 (*n* = 24 fields per reaction; bars specify the median; *, *p* ≤ 0.0001 by Student’s unpaired two-tailed *t*-test). Scale bar equals 5 µm. See also supplemental figure S7 and supplemental table S1.

To ascertain the orientation of the transmembrane state of PEX5, we fused a triple FLAG (3×FLAG) tag to either the N-or C-terminus and repeated the protease protection analysis (Fig. 4B). With the N-terminal fusion, a larger fragment was clipped off by proteinase K, and the resulting truncated form was no longer recognized by FLAG antibodies. On the other hand, the C-terminal fusion showed only a small size shift and retained the FLAG epitope. Thus, PEX5 adopts a transmembrane state with the N-terminus facing the cytosol.

Next, we determined whether the N-terminus of PEX5 always faces the cytosol or first enters the lumen and then emerges. We appended a cleavage site for a lumenal protease to the N-terminus of PEX5, which should cause a size shift of the resulting fusion protein if the N-terminus passes through the lumen (Fig. 4C). Specifically, we used the PTS2 signals from the peroxisomal matrix proteins AGPS and PHYH (**Table S1**), which are cleaved in the lumen by the endogenous protease TYSND1 (Kunze, 2020). The PTS2 motifs were mutated to prevent their targeting function (Dammai and Subramani, 2001) while leaving intact the TYSND1 cleavage sites (**Table S1**). A triple FLAG tag was fused upstream to enhance the size shift, and the conserved cysteine in PEX5 was ablated to prevent any TYSND1-cleaved form from leaving the organelle. The results show that a considerable fraction of the peroxisome-associated population was processed to a smaller form, while no processing occurred after mutating the TYSND1 cleavage sites (Fig. 4D). Thus, the N-terminus of PEX5 must have entered the lumen.

We then assessed whether the TYSND1-cleaved form could still reach the transmembrane state (Fig. 4E). Indeed, the cleaved form of the PHYH fusion was further truncated by addition of proteinase K, similarly to PEX5 with the native N-terminus, indicating that it had reached a transmembrane orientation (lanes 1-4). In contrast, the full-length precursor that co-purified with peroxisomes was completely degraded even at a low concentration of proteinase K, revealing that it resided predominantly on the outside of the organelles. In the absence of TYSND1 cleavage (lanes 5-8), a significant amount of the full-length precursor remained protected from degradation at the low proteinase K concentration. This pool had also reached the transmembrane state, because at a high proteinase K concentration, it was entirely converted into the truncated form indicative of a transmembrane topology. Thus, the N-terminus of PEX5 passes through the peroxisomal lumen before emerging in the cytosol. Notably, this conclusion is consistent with the lumenal localization of the N-terminal domain of PEX14, which provides a strong binding site inside peroxisomes for the pentapeptide import signals.

We next determined whether the entire PEX5 molecule, or just the N-terminus, enters the peroxisomal lumen. We fused the TYSND1 cleavage signal from PHYH to the C-terminus of the receptor, or inserted it inbetween the receptor’s unstructured N-terminal region and the TPR domain (Fig. 4C). In each case, a cleaved fragment was observed in the peroxisome-associated population, similarly to the fusion construct with the TYSND1 cleavage signal positioned at the N-terminus (Fig. 4F). No cleavage was discernible for any of the constructs after mutating the TYSND1 site. The version of each fusion protein with the conserved cysteine intact showed a similar cleavage pattern (**fig. S7E**) and could mediate protein import into peroxisomes (Fig. 4G), indicating that all three fusions were functional (although the N-terminal fusion was less active). Taken together, our results indicate that PEX5 completely enters the peroxisomal lumen during import, and then exits the lumen by a transmembrane intermediate whose N-terminus is in the cytosol.

### A signal for PEX5 export

We next determined the signals in PEX5 that enable exit from the lumen. The amphipathic helices AH1 and AH2 were likely candidates, as they appear to act downstream from docking (Fig. 3E). We ablated the hydrophobic surface of either helix, and examined the topology of the resulting mutants by protease protection (Fig. 5A). The AH2 mutant was fully protected from proteinase K, demonstrating that it resided entirely within the lumen. Thus, this helix is indeed a signal for export of PEX5. In contrast, the AH1 mutant was truncated by the protease, indicating that its N-terminus had emerged in the cytosol, similarly to wild type PEX5 and the C11A version. Thus, both the AH1 and C11A mutants allowed the N-terminus of PEX5 to emerge in the cytosol and were blocked at a later step. These results were confirmed by fusing a triple FLAG tag to the constructs to increase the size shift after protease digestion (Fig. 5B).

**Figure 5.**
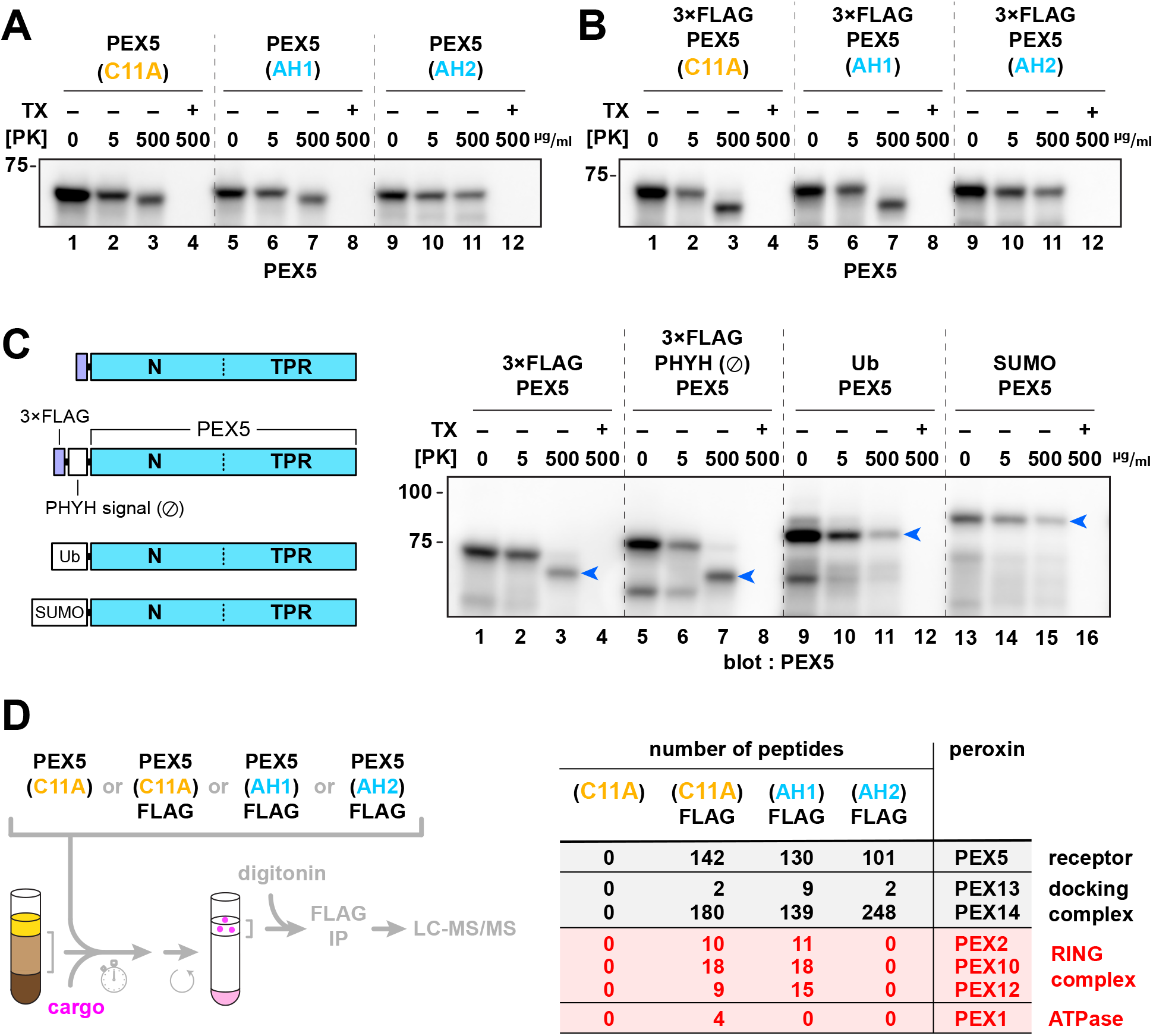
A signal for PEX5 export. (A) *Xenopus* egg extract was incubated with GFP-SKL and PEX5 lacking the conserved cysteine (C11A), or with mutants in which the residues along the hydrophobic face of AH1 or AH2 were mutated to alanine. The peroxisome-associated population was isolated by flotation and treated with different concentrations of proteinase K, with or without Triton X-100 (TX), and immunoblotted for PEX5. Relative molecular weights (kD) are marked on the left. (B) As in (A), except that the mutants were fused at the N-terminus to a triple FLAG (3×FLAG) tag. (C) As in (A), except that the constructs depicted in the scheme on the left were used: 3×FLAG-tagged PEX5; 3×FLAG-tagged PEX5 with an unstructured non-cleavable signal from PHYH; PEX5 fused to ubiquitin (Ub) via a non-hydrolyzable linker; and PEX5 fused to a non-hydrolyzable SUMO. Blue arrows indicate protease-resistant fragments. (D) Extract was incubated with GFP-SKL and the indicated PEX5 variants. Peroxisomes were isolated by flotation, solubilized in digitonin, and the PEX5-associated population was isolated with anti-FLAG beads. Bound proteins were identified by mass spectrometry. The total number of peptides corresponding to the indicated peroxins recovered with each construct is shown on the right. See also supplemental figure S7.

Formation of the transmembrane export intermediate requires the N-terminus of PEX5 to be unstructured. When folded domains (ubiquitin or SUMO) were appended to the N-terminus of PEX5, a considerable fraction of each fusion protein was entirely protected from digestion by proteinase K, and no truncated products were discernible (Fig. 5C). Thus, folded domains at the receptor’s N-terminus block export, in contrast to flexible FLAG tags or unstructured polypeptides (Fig. 5C).

Next we tested whether the retro-translocation intermediates generated with the export-deficient mutants (i.e., AH1, AH2, C11A) interact with the recycling machinery, consisting of the ubiquitin ligase complex and the PEX1/PEX6 ATPase. We fused each mutant to a C-terminal FLAG tag, which does not affect the import activity of wild type PEX5 (**fig. S7F**), and isolated the peroxisome-associated population by flotation. After solubilization in digitonin, interacting proteins were immunoprecipitated with FLAG antibodies. Given the low abundance of the recycling components in *Xenopus* extract (Wühr et al., 2014), we identified interacting proteins by mass spectrometry (Fig. 5D).

With the C11A mutant, all three members of the ubiquitin ligase complex (i.e., PEX2, PEX10, and PEX12) were precipitated. The complex seemed to be considerably enriched, as it is below the limit of detection in the total extract (Wühr et al., 2014). The ATPase subunit PEX1 was also recovered. Given that the C11A mutant cannot be monoubiquitinated and adopts a transmembrane orientation with the N-terminus in the cytosol, the interaction with both the ligase and the ATPase is consistent with this mutant being stalled prior to extraction from peroxisomes. The mutant also prominently precipitated the PEX14 component of the docking complex, and to a lesser extent PEX13. The interaction with the docking complex is consistent with the pentapeptide motifs of PEX5 interacting with the lumenal domain of PEX14. Crucially, none of these peroxins was recovered in the absence of the FLAG tag.

The AH1 mutant also precipitated the ligase complex and the docking components, but not PEX1. Thus, this mutant may be arrested at a different step than C11A, possibly in recruiting the ATPase. Consistent with the AH2 mutant being in the lumen, it did not precipitate any member of the ubiquitin ligase complex or the ATPase, whereas the interaction with the docking components was unaffected. Thus, the observed export defect of the AH2 mutant can be attributed to its inability to engage the ligase, which would in turn preclude the interaction with the ATPase. The AH2 helix is therefore necessary to form the transmembrane intermediate in which the N-terminus is in the cytosol and the C-terminus, including the folded TPR domain, still inside the peroxisomal lumen.

### Unfolding of PEX5 during export

Finally, we asked whether extraction of PEX5 out of the peroxisomal lumen involves unfolding of the C-terminal TPR domain. We first replaced the TPR domain with a nanobody against GFP (Kirchhofer et al., 2010), retaining the entire unstructured N-terminus of PEX5 with all of the features needed for import and recycling, and tested whether the resulting construct would import GFP lacking a peroxisomal targeting signal (Fig. 6A). If the nanobody remains bound to GFP, only a single round of import should occur. On the other hand, if the interaction is broken during each import cycle by unfolding of the nanobody, GFP should accumulate in peroxisomes. The results show that GFP accumulated (Fig. 6B, panel 8, and Fig. 6D). The nanobody construct mediated multiple import cycles, because little GFP accumulation was observed with the non-recyclable C11A mutant of the nanobody fusion (panel 10). Wild type PEX5 was inactive (Fig. 6B, panel 4), although it efficiently imported GFP-SKL (panel 3). We then fused the GFP nanobody to the C-terminus of full-length PEX5 (Fig. 6A). Again, this construct supported multiple rounds of GFP import (Fig. 6C and D), suggesting that both the TPR domain and the nanobody were unfolded during PEX5 export. Because PEX5’s TPR domain fully enters the lumen, it likely remains folded during import to deliver bound cargo into peroxisomes. Unfolding of the TPR domain during export provides a plausible mechanism for how imported cargo is stripped from PEX5 and released inside the lumen.

**Figure 6.**
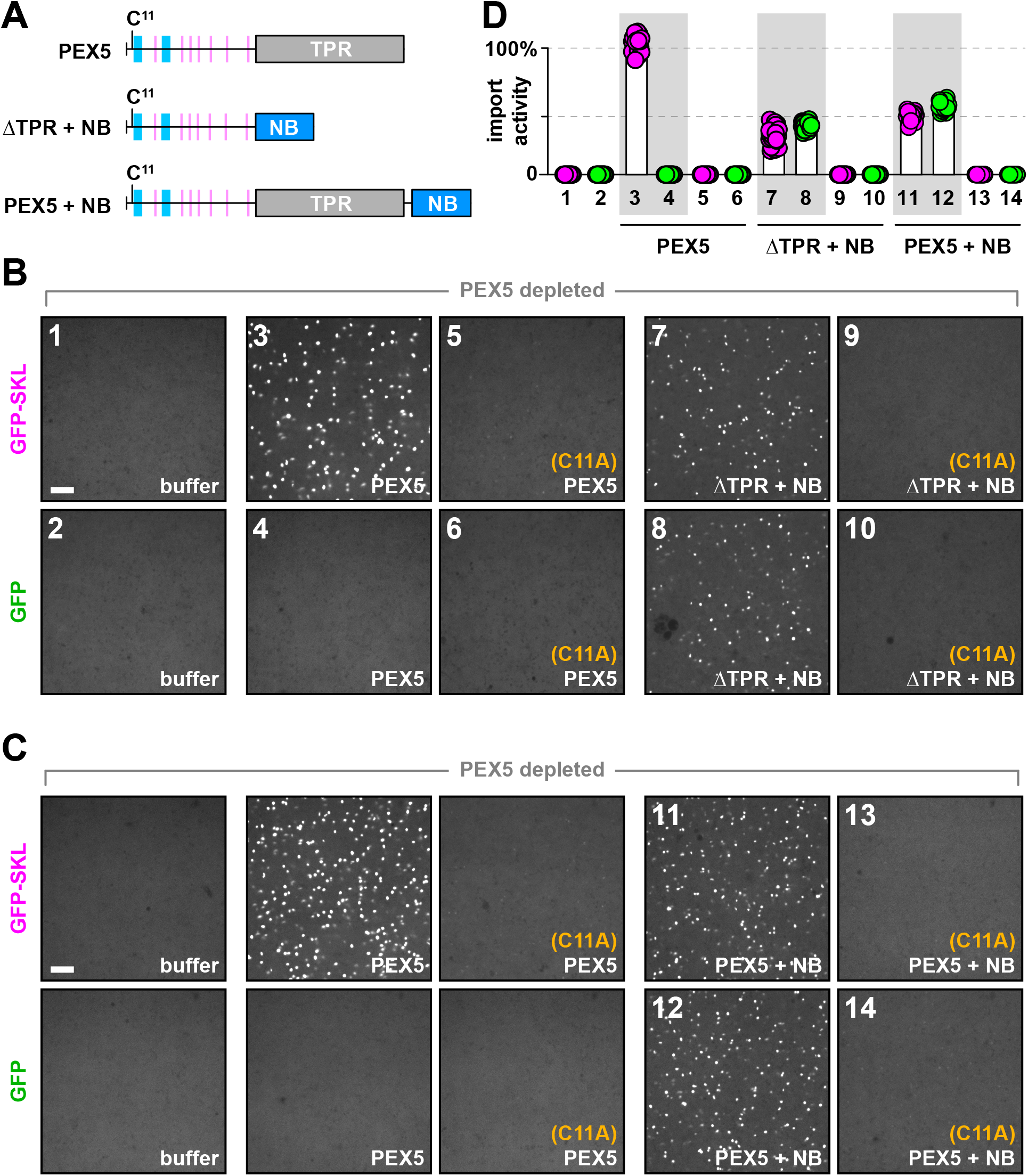
Unfolding of PEX5 during export. (A) Scheme depicting the constructs used to evaluate whether PEX5 is unfolded during its export from peroxisomes. NB denotes a nanobody against GFP that was fused either to the N-terminal unstructured region of PEX5 (ΔTPR + NB) or to the full-length protein (PEX5 + NB). (B) *Xenopus* egg extract was depleted of endogenous PEX5 using beads conjugated to the PEX5-binding domain from PEX14, then supplemented either with wild type PEX5, PEX5 lacking the conserved cysteine (C11A), or PEX5 (ΔTPR + NB) with or without the C11A mutation. Import activity was assessed by incubating with GFP-SKL and imaging the formation of bright puncta by spinning disk confocal microscopy. (C) As in (B), except that the PEX5 + NB construct was used, with or without the C11A mutation. (D) The number of puncta in an imaged field, for the numbered reactions in (B) and (C), was quantified relative to that in the reaction with wild type PEX5 (*n* = 24 fields per reaction; bars specify the median). All scale bars equal 5 µm.

## Discussion

Our results, combined with data in the literature, lead to the following model for how the receptor PEX5 mediates import of peroxisomal matrix proteins (Fig. 7). First, the receptor binds PTS1 matrix proteins in the cytosol through its C-terminal TPR domain (step 1). The cargo-bound receptor is then recruited to peroxisomes using several N-terminal pentapeptide motifs (step 2). Next, PEX5 crosses the membrane into the lumen, taking bound cargo along (step 3). Diffusion of PEX5 and bound cargo back into the cytosol is prevented by the high-affinity interaction between the pentapeptide motifs and the lumenal domain of PEX14. To initiate recycling, the receptor’s unstructured N-terminus emerges in the cytosol by inserting into the ubiquitin ligase complex (step 4). The interaction with the ligase requires a conserved amphipathic helix (AH2) within the unstructured N-terminal region of PEX5. The receptor is then monoubiquitinated at the conserved cysteine (step 5), and completely pulled out of the lumen into the cytosol by the PEX1/PEX6 ATPase (step 6). These steps require a second conserved amphipathic helix (AH1) near the receptor’s N-terminus. Pulling by the ATPase unfolds the C-terminal TPR domain, causing cargo to be released inside the lumen. Finally, the TPR domain refolds in the cytosol and the ubiquitin is removed by deubiquitinating enzymes (step 7), thereby resetting the receptor for a new import cycle.

**Figure 7.**
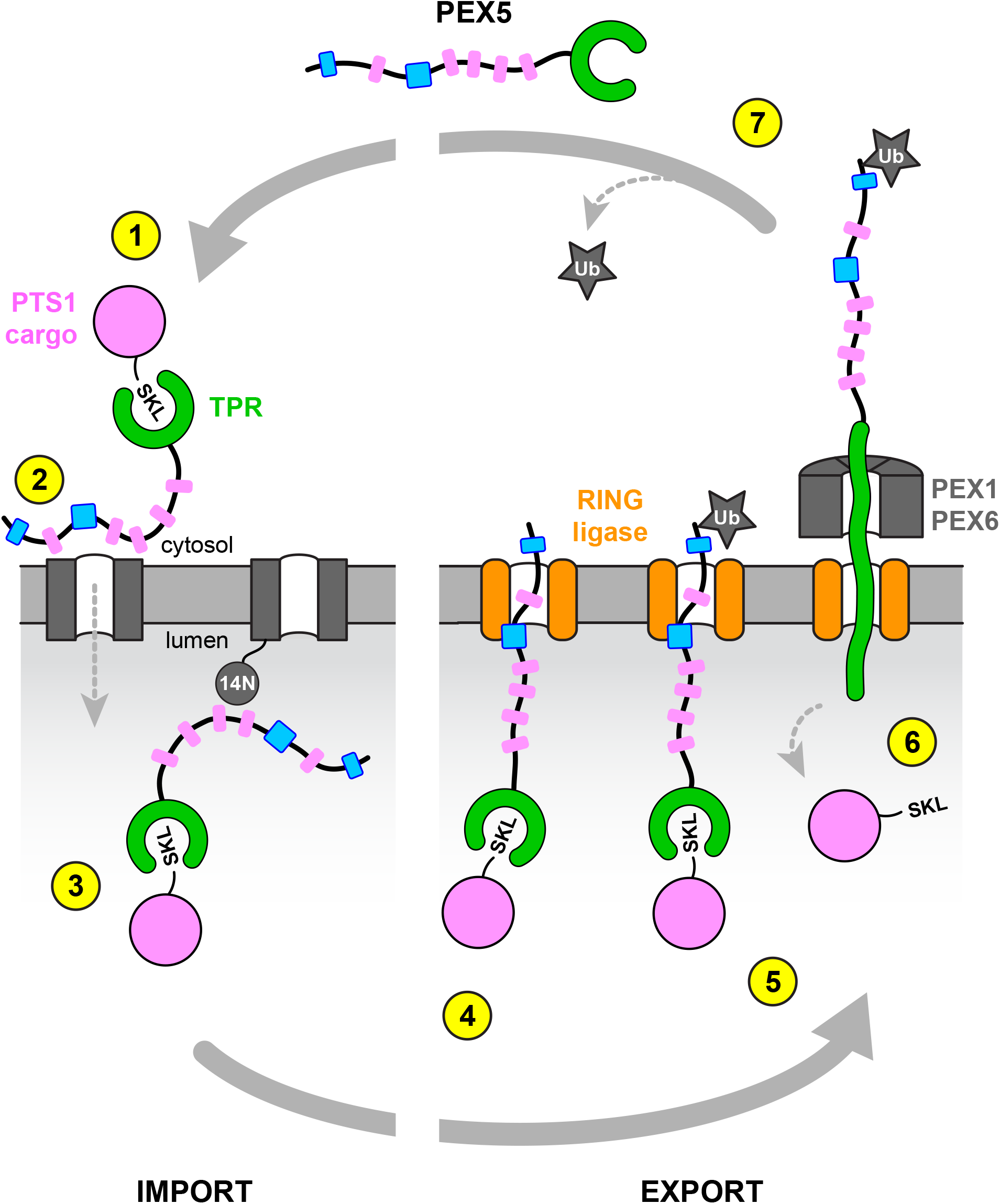
Model of PEX5 shuttling. Step 1: PTS1 cargo binds to the TPR domain of PEX5 in the cytosol. Step 2: Cargo-bound PEX5 is recruited to the docking complex in the peroxisomal membrane, composed of PEX13 and PEX14, using WxxxF/Y pentapeptide motifs (pink). Only the first 5 motifs are shown. Step 3: PEX5 translocates all the way into the lumen along with cargo. Diffusion back into the cytosol is prevented by the high-affinity interaction between the pentapeptide motifs and the lumenal domain of PEX14. Step 4: Export of PEX5 is initiated by binding to the ligase complex through an amphipathic helix (blue). The unstructured N-terminus then inserts into the ligase pore and emerges in the cytosol. Step 5: The conserved cysteine in PEX5 is monoubiquitinated. Step 6: PEX5 is pulled out of the lumen through the ligase pore by the PEX1/PEX6 AAA ATPase. Extraction is accompanied by unfolding of the TPR domain and release of bound cargo. Step 7: The TPR domain refolds in the cytosol and ubiquitin is removed by deubiquitinating enzymes, resetting PEX5 for a subsequent import cycle.

Whether PEX5 moves completely into the peroxisomal lumen has been controversial (Dammai and Subramani, 2001; Erdmann and Schliebs, 2005). Our data support the translocation of the entire PEX5 molecule. It seems less likely that the receptor would be a membrane-embedded component of a translocation channel, as PEX5 has no hydrophobic segments that could insert into the membrane (Emmanouilidis et al., 2016). The only moderately hydrophobic segments required for import are the two amphipathic helices AH1 and AH2, whose absence we show has no effect on PEX5’s ability to traverse the membrane. Complete translocation of cargo-bound receptor across the peroxisomal membrane is also consistent with the known import of folded proteins; the folded TPR domain of PEX5 would simply remain associated with the PTS1 signal of the cargo. PEX5 is the only known import receptor that crosses the membrane. In other systems of protein import into organelles, such as that of the endoplasmic reticulum (ER), mitochondria, or chloroplasts, cargo is recognized by receptors that are either permanently bound to the target membrane or move from the cytosol to the cytosolic surface of the membrane (e.g., the signal recognition particle).

We show that all of the features necessary for PEX5 shuttling are located within the first half of the receptor’s unstructured N-terminal region. The required features comprise the pentapeptide motifs (W0-W4) that mediate peroxisomal recruitment, as well as the conserved cysteine and two amphipathic helices (AH1 and AH2) necessary for recycling. The amino acids located inbetween these features are less conserved, both in terms of their number and identity, and presumably function as linkers that maintain the unstructured nature of the region. The function of the downstream pentapeptide motifs (W5-W7) and amphipathic helices (AH3 and AH4) is unclear. Our experiments show that they are not required for import of a cargo with a canonical PTS1 signal, but they could provide binding sites for proteins that are imported independently of PEX7 or of the TPR domain of PEX5 (Gunkel et al., 2004; Kempiński et al., 2020; Rymer et al., 2018).

How cargo-loaded PEX5 associates with the docking complex and crosses the peroxisomal membrane remains enigmatic. While it is clear that the pentapeptide motifs in PEX5 are required, the motifs interact most strongly with a domain of PEX14 that clearly faces the lumen (this study and Barros-Barbosa et al., 2019). The motifs also bind to a domain in PEX13, which again has been reported to face the lumen (Barros-Barbosa et al., 2019). One possibility is that these domains, particularly that of PEX14, transiently flip across the membrane, capturing PEX5 in the cytosol and then pulling the bound receptor into the lumen (Yamashita et al., 2020). Alternatively, some of the pentapeptide motifs might initially bind weakly to other regions of the docking complex (Otera et al., 2002), and then contact the stronger binding domains in the lumen following translocation. In any case, the tight interaction with the docking complex in the lumen could drive translocation by preventing PEX5 from diffusing back into the cytosol. Avidity might potentiate the interaction, because PEX5 proteins often contain multiple pentapeptide motifs (Emmanouilidis et al., 2016) and the docking complex likely consists of multiple copies of PEX13 and PEX14 (Fransen et al., 2002; Krause et al., 2013; Lill et al., 2020; Oliveira et al., 2002). Sustained import would therefore require unliganded docking components, which could only be generated by continual export of PEX5 from the lumen. The need to overcome avidity and dissolve these strong interactions would explain why PEX5 export is ATP-dependent, whereas import *per se* is not (Miyata and Fujiki, 2005; Oliveira et al., 2003; Platta et al., 2005). This model posits that import and export are coupled, because new receptor molecules can only enter the lumen after the previously imported molecules have been exported and lumenal binding sites have been regenerated.

Our results clarify previously conflicting observations regarding the localization of the receptor’s N-terminus during import. Experiments in cells had suggested that the N-terminus of PEX5 enters the peroxisomal lumen (Dammai and Subramani, 2001), whereas experiments *in vitro* had argued that PEX5 adopts a transmembrane orientation in the peroxisomal membrane with the N-terminus in the cytosol (Gouveia et al., 2003, 2000). We now demonstrate that these observations reflect distinct steps in the shuttling mechanism of PEX5: the N-terminus first enters the lumen during import, and then emerges back in the cytosol to initiate recycling. The transmembrane topology of PEX5 thus corresponds to a populated retro-translocation intermediate.

We recently determined a cryogenic electron microscopy (cryo-EM) structure of the peroxisomal ubiquitin ligase complex (Feng et al., 2022), which can explain the transmembrane topology of the retro-translocation intermediate. The three subunits of the ligase complex (i.e., PEX2, PEX10, and PEX12) co-assemble in the membrane into a channel with an open ∼10-Å pore. The RING finger domains that catalyze receptor ubiquitination are positioned above the pore on the cytosolic side of the complex. Given that the channel does not have a lateral gate open to the membrane, receptors must access the RING domains by inserting their unstructured N-termini into the pore from the lumenal side. The location of the cysteine in PEX5 (position 11 in humans and other vertebrates) suggests that approximately 20-30 N-terminal amino acids would protrude from the pore into the cytosol. This length would be sufficient to expose both the conserved cysteine and the first amphipathic helix (AH1) to the cytosol, and is consistent with the ∼2-kD size shift observed in protease protection experiments. The narrow diameter of the ligase pore would additionally explain why folded domains fused to the receptor’s N-terminus are not tolerated, and why the C-terminal TPR domain has to be unfolded during retro-translocation by the PEX1/PEX6 ATPase. Unfolding of the TPR domain provides a simple mechanism of cargo release, but the involvement of other components might facilitate the process (reviewed in Ma et al., 2011).

We show that the interaction of PEX5 with the ligase complex requires the second amphipathic helix (AH2) in the N-terminal region of PEX5. Given that only the hydrophobic face of this helix is conserved, AH2 might use this surface to bind the lumenal side of the ligase complex and position the receptor’s N-terminus for insertion into the ligase pore. AH2 would thus restrict the length of the N-terminus that is spooled out into the cytosol, thereby localizing the conserved cysteine at the optimal height above the membrane for ubiquitination by the RING domains. Consistent with this model, the size of the segment that stays in the lumen is unaffected by fusing an unstructured segment to the N-terminus of PEX5. The function of the first amphipathic helix (AH1) is less clear. Given its proximity to the conserved cysteine, it could be involved in monoubiquitination. Alternatively, it might help to recruit the PEX1/PEX6 ATPase, because its mutation abolished the interaction in our pull-down/mass spectrometry experiments.

Receptor export is strikingly similar to ER-associated protein degradation (ERAD). In ERAD, multispanning ubiquitin ligases provide a passageway for misfolded ER proteins through the membrane; once on the cytosolic side, these proteins are ubiquitinated and extracted from the membrane by the p97 AAA ATPase (Wu and Rapoport, 2018). Similarly, retro-translocation of PEX5 is initiated by an interaction of the unstructured N-terminus with a multi-spanning ubiquitin ligase complex, followed by receptor ubiquitination in the cytosol and extraction by the PEX1/PEX6 AAA ATPase.

Our results with *Xenopus* egg extract confirm that monoubiquitination is sufficient for the function of PEX5 in peroxisomal matrix protein import, as ubiquitin can be replaced with methylated ubiquitin and the conserved cysteine in PEX5 is the only ubiquitination site required. However, so far, we have been unable to detect a significant percentage of thioester-linked monoubiquitinated PEX5, perhaps because this species is too labile or transient.

The proposed model also explains how PTS2 cargo is imported. The ternary complex, consisting of PEX5, cargo, and the adapter PEX7, would dock to peroxisomes and cross the membrane using the receptor’s pentapeptide motifs. Subsequent export of PEX5 through the ligase pore would strip off both the cargo and PEX7 and leave them behind in the lumen. How PEX7 returns to the cytosol is unclear, as it lacks unstructured regions that could insert into the ligase pore.

## Acknowledgments

We thank Dirk Görlich (Max Planck Institute) for the plasmid encoding *B. distachyon* SUMO, Peiqiang Feng for purification of mouse UBA1, and Yuliia Motrenko for performing the E2 charging assay. We are also grateful to Peiqiang Feng and Yuan Gao for insightful discussions and helpful comments on the manuscript. Fluorescence microscopy was performed at the Nikon Imaging Center at Harvard Medical School with the assistance of Jennifer Waters. Coverslips were prepared for imaging at The Microfluidics Core Facility at Harvard Medical School with the assistance of Calixto Saenz. M.L.S. is a Howard Hughes Medical Institute Fellow of the Helen Hay Whitney Foundation. T.A.R. is a Howard Hughes Medical Institute investigator. This work was supported by a NIGMS grant (R01 GMO52586) to T.A.R.

## Author Contributions

M.L.S. and T.A.R. designed the experiments and wrote the paper; M.L.S. conducted the experiments.

## STAR METHODS

### RESOURCE AVAILABILITY

#### Lead contact

Further information and requests for resources and reagents should be directed to and will be fulfilled by the lead contact, Tom A. Rapoport.

#### Materials availability

All unique reagents generated by this study are available from the lead contact with a completed Materials Transfer Agreement.

#### Data and code availability

- Fluorescence micrographs, immunoblot images, and mass spectrometry data reported in this paper will be shared by the lead contact upon request.
- This paper does not report original code. Fluorescence micrographs and immunoblot images were analyzed using routine functions in ImageJ as described in the relevant STAR Methods sections.
- Any additional information required to reanalyze the data reported in this paper is available from the lead contact upon request.

### EXPERIMENTAL MODEL AND SUBJECT DETAILS

#### X. laevis husbandry

*X. laevis* wild type pigmented African clawed frogs (female, aged 9-20 years) were housed and bred in the Amphibian/Aquatics facility of the Cell Biology Department at Harvard Medical School. Frogs were maintained at 18 °C on a 12-h broad-wavelength light cycle and a biweekly feeding schedule, in water purified by reverse osmosis and conditioned with rock salt approved for aquaculture. Frogs were handled and ovulated strictly within the confines of the facility according to protocols approved by the Harvard Institutional Animal Care and Use Committee.

### METHOD DETAILS

All reagents were from purchased from Millipore-Sigma unless indicated otherwise. All buffers were prepared in ultrapure water. RT denotes room temperature.

#### Plasmid construction

All recombinant proteins were produced as fusions containing N-terminal glutathione-S-transferase (GST), unless indicated otherwise. The sequence coding for each recombinant protein was codon-optimized for *Escherichia coli* and inserted downstream of the GST coding sequence in plasmid pET-28b(+) by Gibson Assembly (from New England BioLabs). A 3C protease cleavage site was incorporated between the GST and the recombinant protein. All constructs were verified by sequencing.

The sequence encoding full-length, wild type PEX5 (designated the long isoform) corresponds to isoform X1 of the *X. laevis* PEX5.S gene (GenBank accession no. XP_018082760.1). The short isoform lacking the PEX7-binding region was generated by removing amino acids 205-244 by PCR and religating the resulting product, and corresponds to isoform X3 (GenBank accession no. XP_018082765.1). Additional deletions and mutations of PEX5 mentioned in the text were generated by a similar strategy. Note that PEX5 constructs shown in Figures 1 and 2 were generated in the long isoform background, whereas all other figures used the short isoform. FLAG-tagged constructs were assembled by fusing the sequence encoding the amino acids DYKDDDDK (1×FLAG) or DYKDHDGDYKDHDIDYKDDDDK (3×FLAG) to either the N-or C-terminus of wild type PEX5 or the mutants indicated in the text, separated by a (GGS)_2_ linker.

PEX5 fused to the PTS2 import signal from AGPS (alkylglycerone phosphate synthase) was generated as follows. The sequence encoding residues 1-51 of *X. laevis* AGPS.L (GenBank accession no. AAH76825.1) was appended to the N-terminus of the short isoform of PEX5, separated by a (GGS)_2_ linker. The PTS2 motif was inactivated by mutating Leu18 to Arg within the import signal, and a 3×FLAG tag was then inserted upstream of the resulting construct, separated by a (GGS)_2_ linker. To inactivate the TYSND1 cleavage site, amino acids 33-40 within the import signal were replaced with the sequence KLGSSGELS. Where indicated in the text, the conserved cysteine (Cys11) in PEX5 was also mutated to Ala.

PEX5 fused at the N-terminus to the PTS2 import signal from PHYH (phytanyl-CoA hydroxylase) was constructed similarly, except that residues 1-40 of *X. laevis* PHYH.S (GenBank accession no. XP_018109771.1) were used, the PTS2 motif was inactivated by mutating Leu16 within the import signal to Arg, and the TYSND1 site was inactivated by converting residues 25-33 within the import signal to the sequence DGSKLGGGS. For the C-terminal constructs, the PHYH import signal was fused to the C-terminus of the short PEX5 isoform separated by a GSG linker, followed by a spacer consisting of the amino acids GGSGGSMAMRPELGPEDPEAQGGSGGSM, and finally a 3×FLAG tag. For constructs containing the PHYH import signal in the middle of PEX5, the import signal was inserted between residues Ser264 and Ile265 of the short isoform, flanked by GGS linkers.

SUMO-tagged PEX5 was generated by fusing PEX5 directly downstream of the coding sequence of *Brachypodium distachyon* SUMO, which contained three point mutations (i.e., T61K, D68K, and Q76R) to prevent cleavage by endogenous SUMO protease in egg extract (Vera Rodriguez et al., 2019). Ubiquitin-tagged PEX5 was assembled by fusing PEX5 directly downstream of the coding sequence for *X. laevis* UBC.L (GenBank accession no. NP_001080865.1). To prevent removal of the ubiquitin moiety by deubiquitinating enzymes in egg extract, the terminal glycine (Gly76) in ubiquitin was converted to valine. PEX5 constructs fused to a GFP nanobody were assembled by appending the sequence encoding the GFP enhancer nanobody (Kirchhofer et al., 2010) downstream of the full-length short isoform of PEX5, separated by (GGS)_4_ linker, or downstream of the N-terminal region of the short isoform (residues 1-268), separated by a (GGS)_2_ linker.

GFP-SKL was assembled by fusing the sequence encoding the SKL tripeptide, preceded by a glycine, to the coding sequence for mEGFP (Wang et al., 2016). Plasmids encoding mScarlet-SKL, mCherry-SKL, and the *X. laevis* PEX14 N-terminal domain, as well as the purification of the corresponding proteins, were described previously (Romano et al., 2019). The fragment encoding the C-terminal region of PEX14, used for antibody production, corresponds to residues 226-363 of *X. laevis* PEX14.L (GenBank accession no. XP_018081136.1). The sequence encoding UBCH5C (UBE2D3) corresponds to *X. laevis* UBE2D3.L (GenBank accession no. NP_001084434.1). The plasmid encoding mouse UBA1 was acquired from Addgene (no. 32534) and was described previously (Carvalho et al., 2012).

#### Production of recombinant proteins

*E. coli* BL21 Rosetta 2(DE3) cells transformed with the desired plasmid were cultured in baffled flasks on an orbital shaker at 37 °C in 2×YT medium containing 50 µg/ml kanamycin and 34 µg/ml chloramphenicol. When the OD reached 0.6, the cultures were cooled on ice for 30 min, then supplemented with isopropyl β-D-1-thiogalactopyranoside (IPTG) to 1 mM and incubated with shaking at 16 °C for an additional 16 h. Cells were collected by centrifugation at 4000 × *g* for 20 min at 4 °C, then resuspended in purification buffer (50 mM Tris pH 7.8 at RT, 150 mM NaCl, 1 mM EDTA, 1 mM DTT) supplemented with a protease inhibitor cocktail (from Roche, no. 11873580001) according to the manufacturer’s instructions. The cell suspension was passed once through an Avestin EmulsiFlex-C3 high-pressure homogenizer and immediately supplemented with phenylmethylsulfonyl fluoride (PMSF) to 1 mM. The resulting lysate was clarified by centrifugation at 40,000 × *g* for 40 min at 4 °C, and the supernatant was then incubated with glutathione sepharose 4B for 30 min at RT with inversion. Resin was recovered by centrifugation at 500 × *g* for 5 min at 4 °C, and washed extensively in purification buffer, followed by purification buffer supplemented with NaCl to 600 mM, and finally with additional purification buffer. Recombinant protein was eluted by resuspending the washed resin in purification buffer and incubating overnight at 4 °C with homemade GST-tagged 3C protease and gentle agitation. Released protein was collected by draining the resin on a drip column, concentrated on an appropriate Amicon centrifugal concentrator at 4 °C, and gel filtered into XBHS buffer (XB buffer supplemented with HEPES pH 7.8 at RT to 40 mM and 250 mM sucrose) on either a Superdex 75 or 200 Increase 10/300 GL column (from Cytiva). Peak fractions were pooled, concentrated, and snap-frozen in liquid nitrogen in single-use aliquots. All proteins were stored at −80 °C.

For purification of UBCH5C, the GST fusion protein bound to glutathione sepharose resin was washed into 50 mM HEPES•NaOH (pH 7.0 at 4 °C), 150 mM NaCl, 1 mM EDTA, and 1 mM DTT before digestion with 3C protease. All subsequent steps were performed at pH 7.0.

For purification of the C-terminal region of PEX14 for antibody production, the GST fusion protein was eluted from the glutathione sepharose resion using reduced L-glutathione according to the manufacturer’s instructions, concentrated, and gel filtered on a Superdex200 26/600 PG column (from Cytiva) into 25 mM HEPES•KOH (pH 7.7 at RT) and 150 mM KCl. Peak fractions were pooled, concentrated, and supplemented with 250 mM sucrose and 1 mM DTT before snap freezing.

For purification of mouse UBA1, cells were suspended in lysis buffer (25 mM HEPES•NaOH pH 7.4 at RT, 400 mM NaCl, 0.2 mM TCEP (tris(2-carboxyethyl)phosphine), and 50 mM imidazole) in the presence of a protease inhibitor cocktail (from Roche), and lysed by sonication. The clarified lysate was incubated with nickel-charged NTA (nitriloacetic acid) resin for 30 min at 4 °C with agitation, and washed sequentially with lysis buffer, then with lysis buffer supplemented with imidazole to 75 mM. Bound protein was eluted with lysis buffer supplemented with imidazole to 400 mM, mixed with thrombin, and dialyzed overnight at 4 °C into LS buffer (25 mM HEPES•NaOH pH 7.4 at RT, 50 mM NaCl, and 0.2 mM TCEP). Released protein was applied to a Mono Q anion-exchange column (from Cytiva), washed in LS buffer, and eluted using a linear gradient from 50 mM to 1 M NaCl in LS buffer. Peak fractions were pooled, concentrated, and desalted on a Superdex 200 10/300 GL column (from Cytiva) into 25 mM HEPES•NaOH (pH 7.4 at RT), 150 mM NaCl, 0.2 mM TCEP, and 5 % w/v glycerol. Peak fractions were pooled, concentrated, and snap frozen as above.

#### Protein methylation

Primary amine groups on PEX5 were blocked by reductive methylation according to an estalished procedure (Tan 2014). Briefly, 500 µl of a 1 mg/ml solution of recombinant PEX5, dissolved in 50 mM HEPES (pH 7.7 at RT) and 250 mM NaCl, were mixed with 10 µl of 1 M dimethylamine borane (DMAB; from Sigma, no. 180238) and 20 µl of 1 M formaldehyde (from ThermoFisher, no. 28906) and incubated with agitation in the dark at 4 °C. After 2 h, the reaction was supplemented with an additional 10 µl of 1 M DMAB and 20 µl of 1 M formaldehyde, and incubated for an additional 2 h. Finally, the reaction was supplemented with 5 µl of 1 M DMAB and incubated overnight at 4 °C. The following day, reactions were quenched by adding 40 µl of 1 M glycine and 3 µl of 1 M DTT, and incubating for 1 h on ice. The methylated protein was then desalted into XBHS on a Superdex 200 10/300 GL column, and peak fractions were pooled, concentrated, and snap-frozen in liquid nitrogen.

#### Preparation of egg extract

Interphase-locked crude extract lacking filamentous actin was prepared as previously described (Romano et al., 2019), with the following modifications. Eggs were dejellied in Marc’s modified Ringer’s (MMR) solution supplemented with 20 g/L cysteine, 25 mM NaOH, and 50 mM sucrose, and then washed into XB buffer (10 mM Hepes•KOH pH 7.7 at RT, 0.1 M KCl, 1 mM MgCl_2_, 0.1 mM CaCl_2_) supplemented with 50 mM sucrose. Eggs were transferred to 5-ml open-top thinwall tubes (from Beckman, no. 344057) and compacted by centrifugation at RT, first at 350 × *g* for 1 min and then at 1200 × *g* for 40 sec. Excess buffer was removed by aspiration, and a 1000× concentrated cocktail in dimethylsulfoxide (DMSO) was added to yield 10 µg/ml each of pepstatin A, leupeptin, and chymostatin; 10 µg/ml cytochalasin D; and 100 µg/ml cycloheximide. The packed eggs were then disrupted by centrifugation in a swinging bucket rotor at 12,100 × *g* for 15 min at 16 °C. The resulting crude extract was immediately supplemented with protease inhibitors, cytochalasin D, and cycloheximide, and stored on ice until needed.

#### Isolation of membranes from egg extract

Membranes were isolated by flotation through a discontinuous sucrose gradient as previously described (Romano et al., 2019), with the following modifications. For protease protection experiments, 200 µl of crude extract were gently mixed with 400 µl of cold XBH buffer containing 75 % w/v sucrose. 500 µl of the resulting suspension were transferred using a wide bore pipet tip to a 2-ml open-top thinwall tube (from Beckman), and overlaid first with 1.5 ml of cold XBH buffer containing 50 % w/v sucrose, and then with 0.1 ml of cold XBH buffer containing 8 % w/v sucrose. Tubes were centrifuged in a swinging bucket rotor at 220,000 × *g* for 60 min at 4 °C. The membrane band that accumulated at the top of the gradient was gently aspirated using a wide bore tip and transferred to a microfuge tube on ice. Membranes were diluted in 1 ml cold XBHS buffer, collected by centrifugation at 18,000 × *g* for 5 min at 4 °C, and washed again with 1 ml of cold XBHS. The resulting pellet was resuspended in 100 µl of cold XBHS by gently flicking the tube until a homogeneous suspension was obtained.

For mass spectrometry, 500 µl of extract were mixed with 1 ml of cold XBH buffer containing 75 % w/v sucrose. 1 ml of the resulting suspension was transferred to a 5-ml open-top thinwall tube (from Beckman), and overlaid first with 3.8 ml of cold XBH buffer containing 50 % w/v sucrose, and then with 0.2 ml of cold XBH buffer containing 8 % w/v sucrose. Tubes were centrifuged and membranes were harvested as described above. The final membrane pellet was resuspended in 400 µl of cold XBHS. To ensure enough material, each sample was prepared in duplicate and the resulting membrane suspensions were pooled to yield a final volume of 1 ml.

#### Protease protection assay

Proteinase K (from Sigma, no. P6556) was dissolved to 5 mg/ml in XBHS and snap-frozen in single-use aliquots. For each reaction, 18 µl of a membrane suspension (prepared as described above) were mixed with 2 µl of a 10× concentrated solution of proteinase K in XBHS or with XBHS buffer alone. For experiments involving membrane permeabilization, reactions were additionally supplemented with 1 µl of a 10 % w/v solution of high-purity Triton X-100 in water (from Thermo, no. 85111) or with water alone. Reactions were incubated for 1 h on ice or at 24 °C, then quenched by adding 0.5 µl of 400 mM PMSF (in ethanol) and incubating for an additional 10 min at RT. Quenched reactions were diluted 3-fold in boiling Laemmli buffer and heated for 10 min at 98 °C.

#### FLAG immunoprecipitation and mass spectrometry

Membrane suspensions (1 ml) were diluted in XBHS buffer (4 ml) and supplemented with 1 % w/v high-purity digitonin (from Millipore-Sigma, no. 300410) and a protease inhibitor cocktail (from Roche, no. 11836170001) according to the manufacturer’s instructions. The solubilisate was clarified by centrifugation at 21,000 × *g* for 20 min at 4 °C, and the resulting supernatant fraction was then incubated with 300 µl of anti-FLAG M2 affinity gel (from Millipore-Sigma, no. A2220) for 3 h at 4 °C with inversion. The resin was washed with excess XBHS containing 0.1 % w/v digitonin, and bound protein was then eluted with 1 ml of 0.4 mg/ml 3×FLAG peptide (from Bimake, no. B23112) in XBHS containing 0.1 % w/v digitonin. Eluates were concentrated on an Amicon centrifugal concentrator (10-kD MWCO), then heated in Laemmli buffer for 20 min at 60 °C and resolved on a 4-20 % TGX gel (from Bio-Rad). The gel was fixed overnight in 50 % v/v methanol and 5 % v/v acetic acid, then briefly stained with coomassie, and washed multiple times in ultrapure water. Lanes were cut into three sections and submitted to the Taplin Mass Spectrometry Facility at Harvard Medical School for proteomic analysis by LC-MS/MS.

#### Electrophoresis and immunoblotting

Samples were dissolved in Laemmli buffer (50 mM Tris•HCl pH 6.8, 10 % v/v glycerol, 2 % w/v SDS, and 0.01 % bromophenol blue), supplemented with or without 700 mM beta-mercaptoethanol (BME) as indicated in the text, and electrophoretically resolved under denaturing conditions on 4-20 % TGX or single percentage precast polyacrylamide gels (from Bio-Rad). Resolved proteins were transferred to Immun-Blot PVDF membranes (from Bio-Rad, no. 13709A06) at 25 V for 12 h at RT in 25 mM Tris, 192 mM glycine, and 10% v/v methanol. Membranes were blotted in TBST (20 mM Tris•HCl pH 7.5, 150 mM NaCl, and 0.01 % v/v Tween 20), supplemented with 3 % w/v fat-free dried milk, with polyclonal antibodies against full-length *X. laevis* PEX5 or the N-terminal domain of *X. laevis* PEX14 (both from Romano et al., 2019); the C-terminal unstructured region of *X. laevis* PEX14 (this study); or against the FLAG epitope (from Sigma, no. F7425). Blots were developed by chemiluminescence and imaged on an Amersham Imager 600 instrument.

#### Antibody production

Antibodies against the C-terminus of *X. laevis* PEX14 were raised in rabbits by Thermo Fisher Scientific, using the purified fragment fused to GST, and affinity purified from serum as previously described (Romano et al., 2019) using the antigen immobilized on beads.

#### Peroxisomal import reactions

For a typical experiment, crude extract was first manipulated either by adding reagents or removing endogenous components. The effect on import activity was then assessed by adding a concentrated solution of a fluorescent cargo (as indicated in the text) to 0.5 µM and incubating at 24 °C for 1 h. Reactions were then immediately prepared for imaging as decribed below. The E1 activating enzyme inhibitors MLN-7243 (no. S7109) and MLN-4924 (no. S8341) were from SelleckChem. The deubiquitinating enzyme (DUB) inhibitors ubiquitin vinyl sulfone (Ub-VS; no. U-202) and ubiquitin aldehyde (Ubal; no. U-201) were from Boston Biochem, and PR-619 was from Sigma (no. SML-0430). Mock-treated control reactions received the same volume of solvent or buffer alone.

To deplete endogenous PEX5, crude extracts were tumbled for 30 min at RT with agarose beads covalently conjugated either to polyclonal antibodies against full-length PEX5 or to the high-affinity PEX5-binding domain from PEX14, as previously described (Romano et al., 2019). Import activity of PEX5 mutants was evaluated by supplementing PEX5-depleted extracts with recombinant purified proteins to 1 µM, or with XBHS alone, prior to adding fluorescent cargo.

To deplete endogenous free ubiquitin, extracts were first pre-warmed for 30 min at 24 °C, and then incubated for an additional 1 h at 24 °C with 25 µM UbVS. The ability of ubiquitin or PEX5 mutants to support peroxisomal import was assessed by supplementing the DUB-inhibited extracts with 100 µM recombinant ubiquitin (wild type or mutant) and 0.5 µM recombinant PEX5 (wild type or mutant), prior to adding fluorescent cargo. Wild type and methylated human ubiquitin, and all ubiquitin mutants indicated in the text, were purchased from Boston BioChem and were stored in single-use aliquots at −80 °C.

#### Microscopy

Prior to imaging, 9 µl of an import reaction were sandwiched between two no. 1.5 square glass coverslips (22×22 mm; from VWR, no. 48366-227) that had been passivated with 20-kD polyethylene glycol as described previously (Romano et al., 2019). The sandwich was mounted on a 25 × 75 mm aluminum slide and sealed with VALAP (composed of equal weights of vaseline, lanoline, and paraffin). Reactions were imaged immediately on a spinning disk confocal platform comprised of a Nikon ECLIPSE Ti inverted microscope and a Yokagawa CSU-X1 spinning disk confocal scanner with Spectral Applied Research Aurora Borealis modification. Samples were illuminated by 100 mW solid state 488-nm (Coherent) or 561-nm (Cobolt) lasers selected by an acousto-optic tunable filter (AOTF; from Gooch & Housego). Fluorescence was collected through a 100× 1.45 NA Plan Apochromat Lambda oil immersion objective and a Di01-T405/488/568/647 dichroic (Semrock), using either a green emission ET525/50m (525 ± 25 nm) or a red emission ET620/60m (620 ± 30 nm) single-band bandpass filter (Chroma), respectively. Images were acquired in Metamorph software (64-bit, v. 7.10.3.279) using a Hamamatsu ORCA-R2 cooled CCD camera with 2×2 binning, a high-precision 12-bit digitizer, and analog gain set to 1. For each sample, multiple positions arrayed around the center of the coverslip were imaged using a custom Metamorph script and a Prior Proscan II motorized stage operating at 20 % of the maximum speed (to avoid shaking the extract). To minimize photobleaching and phototoxicity from repeated illumination, positions were spaced 500 µm apart, which corresponds to a distance equal to 5 times the size of an acquired field. Focus was maintained at a preset distance from the coverslip by the Nikon Perfect Focus System (PFS). Imaging parameters were chosen to maximize the SNR while avoiding saturation. All samples intended to be compared were imaged consecutively under identical acquisition settings.

### QUANTIFICATION AND STATISTICAL ANALYSIS

#### Image analysis

All analysis was performed in ImageJ (Schneider et al., 2012) on the original, unmodified image data using custom-written scripts available on request. Images were first manually curated, and any that did not satisfy the following criteria were omitted from analysis: any image that was not properly focused; any image in which relevant features were not stationary; any image containing aggregated extract or other aberrant objects; any image with air bubbles whose combined area exceeded 10 % of the total imaged area; or any image whose total median intensity differed by more than 25 % from the total median intensity of all images from that sample.

Import activity was calculated as the number of peroxisomes visible in an imaged field, and expressed as a percentage of the median number of peroxisomes visible in a control cohort. Note that peroxisomes become visible as diffraction-limited puncta only when the cumulative intensity of imported fluorescent cargo, which we previously showed increases linearly with time (Romano et al., 2019), exceeds the average intensity of unimported cargo outside the organelle. Also note that egg extract is a homogeneous suspension, so that the density of organelles in a particular field is representative, within error, of the density of those organelles in the entire sample. To identify peroxisomes, images were first smoothed by replacing each pixel with the average intensity of all pixels in its 3×3 neighborhood. Local maxima were then identified, such that the difference between each maximum and the background exceeded some tolerance value (see below). Maxima located along the edge of the image were excluded.

The tolerance value was calculated as follows. Images from a positive control reaction (in which import activity is maximal) were first thresholded using the MaxEntropy algorithm in ImageJ to identify puncta that were in-focus, and thus bright, and exclude out-of-focus puncta. The identified compartments were then masked, and the standard deviation of the fluorescence intensity in the remainder of each image was calculated. A value equal to twice the average standard deviation determined from all images in the sample was used as the tolerance.

#### Data plotting and statistical analysis

GraphPad Prism (v. 9.3.1) was used to plot data and perform statistical tests.

#### Image processing for publication

All fluorescence micrographs intended to be compared were first normalized in ImageJ by average background intensity, and then linearly contrast stretched to the same bit range. Digitized images of immunoblots and stained gels were linearly contrast stretched to reveal relevant bands while avoiding clipping of background. Helical wheel diagrams were assembled using HeliQuest (Gautier et al., 2008). Figures were assembled for publication in Adobe Illustrator (v. 25.4.1).

## SUPPLEMENTAL INFORMATION

Supplemental information includes:

Supplemental figures S1-S7

Supplemental Table S1. Supplemental Table S1. Cleavable PTS2 import signals used in this study (related to Figure 4)

