## Supplemental Information for "Mechanism of PEX5-mediated protein import into peroxisomes"

### Supplemental Figure S1

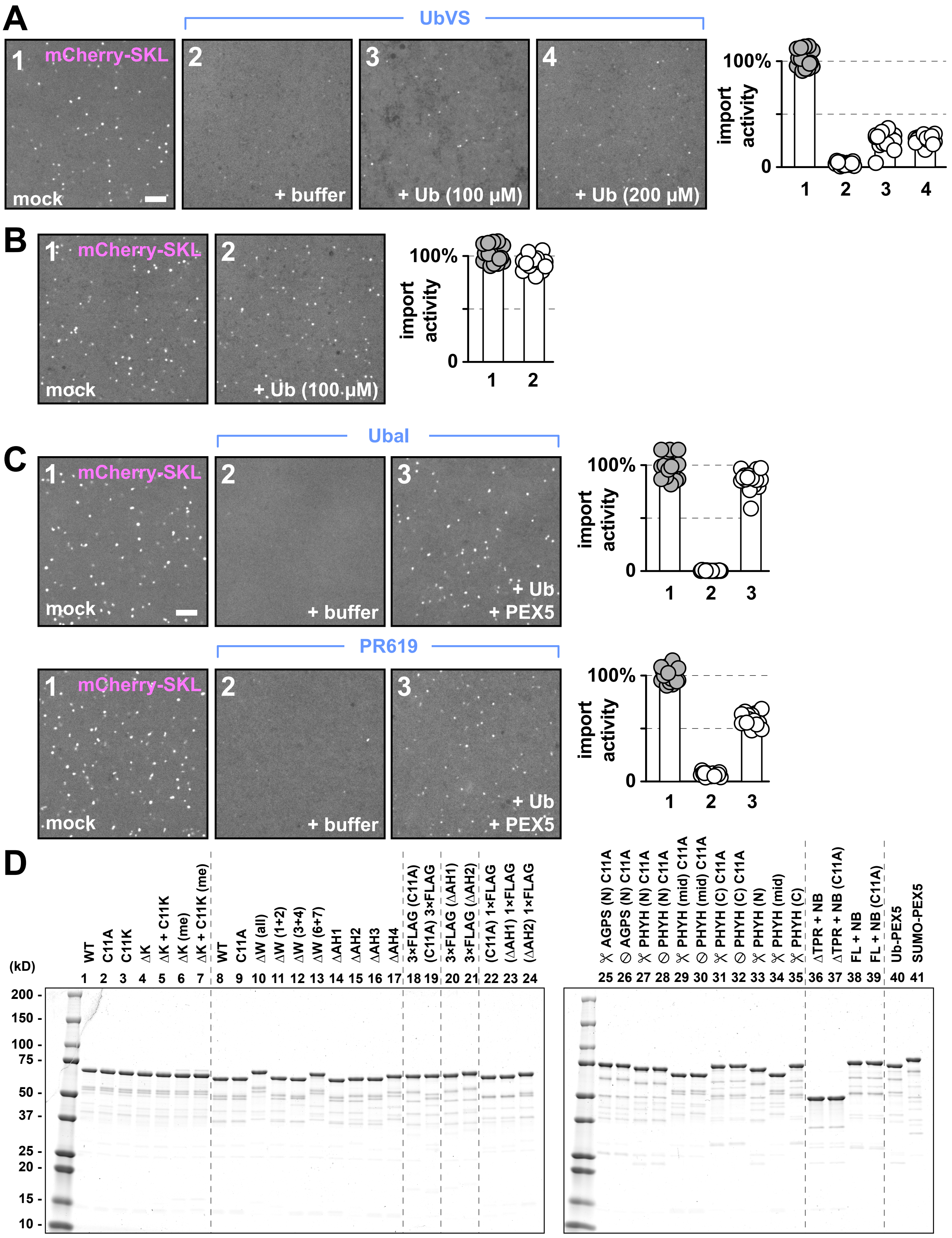

**Supplemental Figure S1. Restoration of peroxisomal protein import in DUB-inhibited extract by ubiquitin and PEX5 (related to Figure 1)**

(A) *Xenopus* egg extract was treated with buffer (mock) or ubiquitin vinyl sulfone (UbVS), and the UbVS-treated reactions were then supplemented with buffer or wild type ubiquitin at the indicated concentrations. Import activity was assessed by incubating the reactions with mCherry-SKL and imaging the formation of bright puncta on a spinning disk confocal microscope. The number of puncta in an imaged field was quantified relative to that in the mock reaction ($n$ = 9 fields per reaction; bars specify the median).

(B) Extract was incubated with mCherry-SKL in the presence of buffer (mock) or 100 μM wild type ubiquitin, and import activity was quantified as above ($n$ = 9 fields per reaction).

(C) As in (A), except that extract was pre-treated with the DUB inhibitors ubiquitin aldehyde (Ubal) or PR-619, a small molecule, then supplemented with buffer or ubiquitin together with PEX5. Import activity was quantified as above ($n$ = 9 fields per reaction). All scale bars equal 5 μm.

(D) Coomassie-stained gels of select purified recombinant PEX5 proteins used in this study. Lanes 1-7 correspond to proteins in Figs. 1 and 2; lanes 8-17 to Fig. 3C-E; lanes 18-19 to Fig. 4B; lanes 20-21 to Fig. 5B; lanes 22-24 Fig. 5D; lanes 25-35 Fig. 4C-G; lanes 36-39 Fig. 6; and lanes 40-41 Fig. 5C.

### Supplemental Figure S2

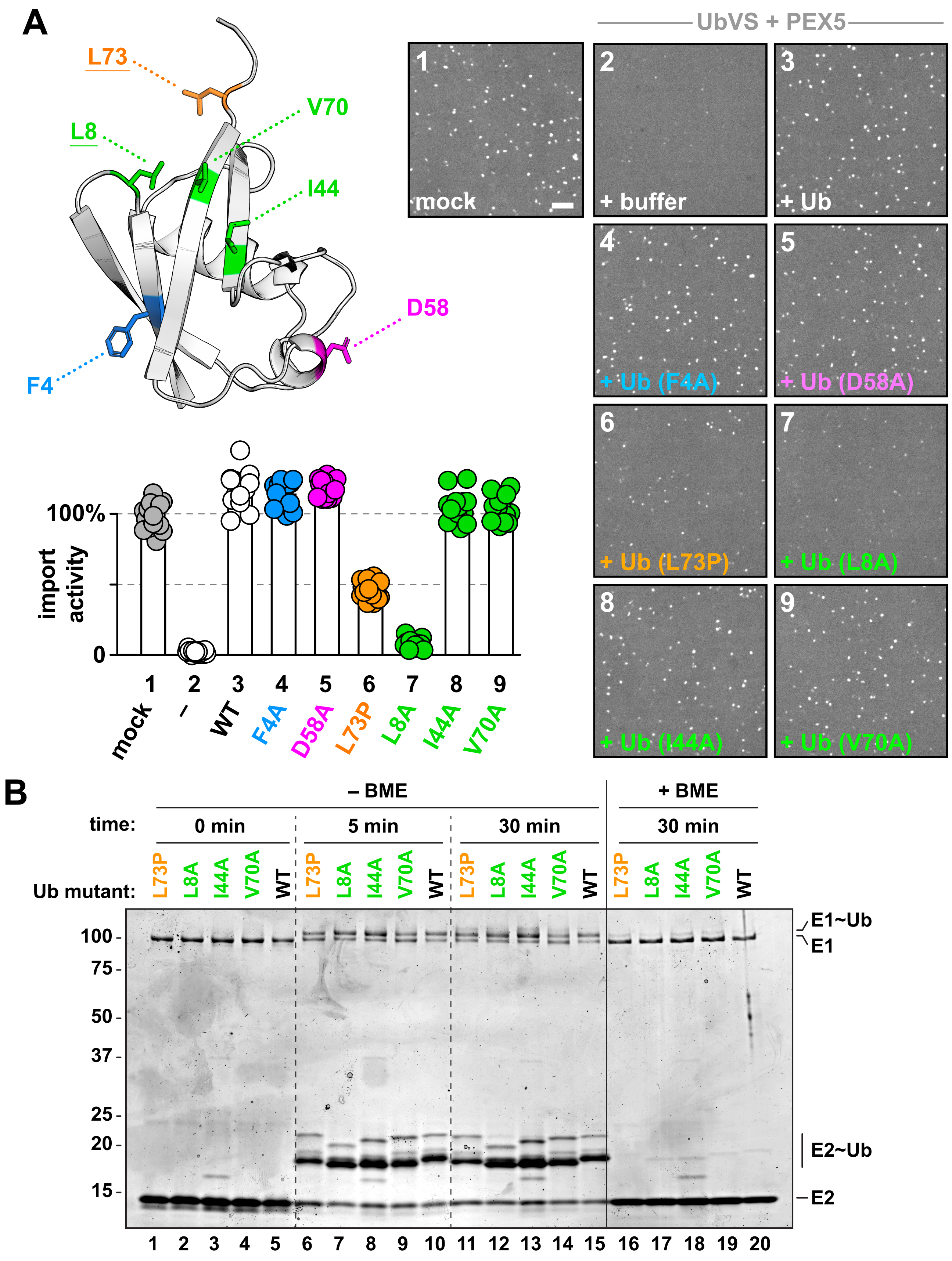

**Supplemental Figure S2. Ubiquitin surface residues required for peroxisomal protein import (related to Figure 2)**

(A) Structural model of ubiquitin (PDB: 1UBQ), depicting key surface residues that were mutated to evaluate their role in peroxisomal import. Mutants lacking the underlined residues did not support import. On the right, the ubiquitin mutants were used in peroxisomal import assays. *Xenopus* egg extract was treated with buffer (mock) or ubiquitin vinyl sulfone (UbVS), and the UbVS-treated reactions were then supplemented with buffer or wild type PEX5 together with the indicated Ub mutants. Import activity was assessed by incubating the reactions with mCherry-SKL and imaging the formation of bright puncta on a spinning disk confocal microscope. The number of puncta in an imaged field was quantified relative to that in the mock reaction ($n$ = 18 fields per reaction; bars specify the median). Scale bar equals 5 µm.
(B) Purified mouse UBA1 (E1), *X. laevis* UBCH5C (E2), and wild type human Ub (WT) or the indicated Ub mutants, were incubated in the presence of ATP and $Mg^{2+}$ for the times shown. Reactions were quenched in SDS-sample buffer with or without the reducing agent β-mercaptoethanol (BME). E1~Ub and E2~Ub denote thioester-linked adducts between Ub and either the E1 or E2, respectively.

### Supplemental Figure S3

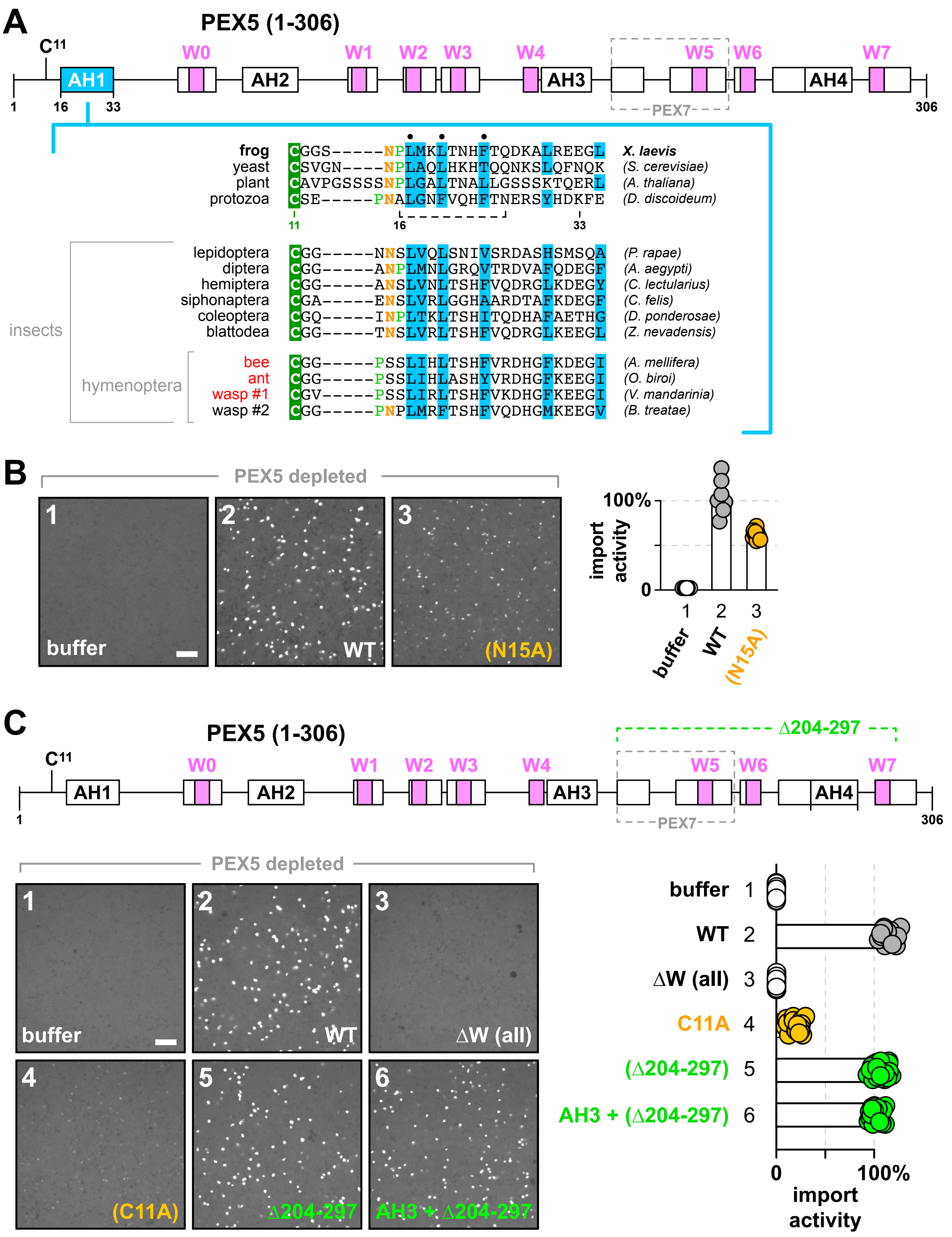

**Supplemental Figure S3. Role of Pex5's conserved asparagine (N15) and amphipathic helices AH3 and AH4 on peroxisomal protein import (related to Figure 3)**

(A) Diagram of the N-terminal unstructured region of *X. laevis* PEX5, showing the locations of predicted α-helices (boxes) and key residues. Amphipathic helices are labeled AH1-AH4; WxxxF/Y pentapeptide motifs are labeled W0-W7. The PEX7-binding region is enclosed by a gray dashed line. Sequence alignments underneath correspond to the region between the conserved cysteine and the C-terminal end of the first amphipathic helix (AH1) in PEX5 homologs from the indicated organisms.

(B) *Xenopus* egg extract was depleted of endogenous PEX5 using beads conjugated to the PEX5-binding domain from PEX14, then supplemented either with buffer, wild type PEX5 (WT), or a PEX5 mutant in which Asn15 was converted to alanine (N15A). Import activity was assessed by incubating the reactions with mCherry-SKL and imaging the formation of bright puncta on a spinning disk confocal microscope. The number of puncta in an imaged field was quantified relative to that in the reaction with wild type PEX5 ($n$ = 9 fields per reaction; bars specify the median).

(C) As in (B). C11A refers to a PEX5 mutant in which Cys11 was mutated to Ala; ΔW (all) refers to a mutant in which all WxxxF/Y motifs were mutated to AxxxA; Δ204-297 lacks the indicated residues (shown in the diagram on top); and AH3 + Δ204-297 additionally has the residues along the hydrophobic face of AH3 mutated to alanines. All scale bars equal 5 μm.

### Supplemental Figure S4

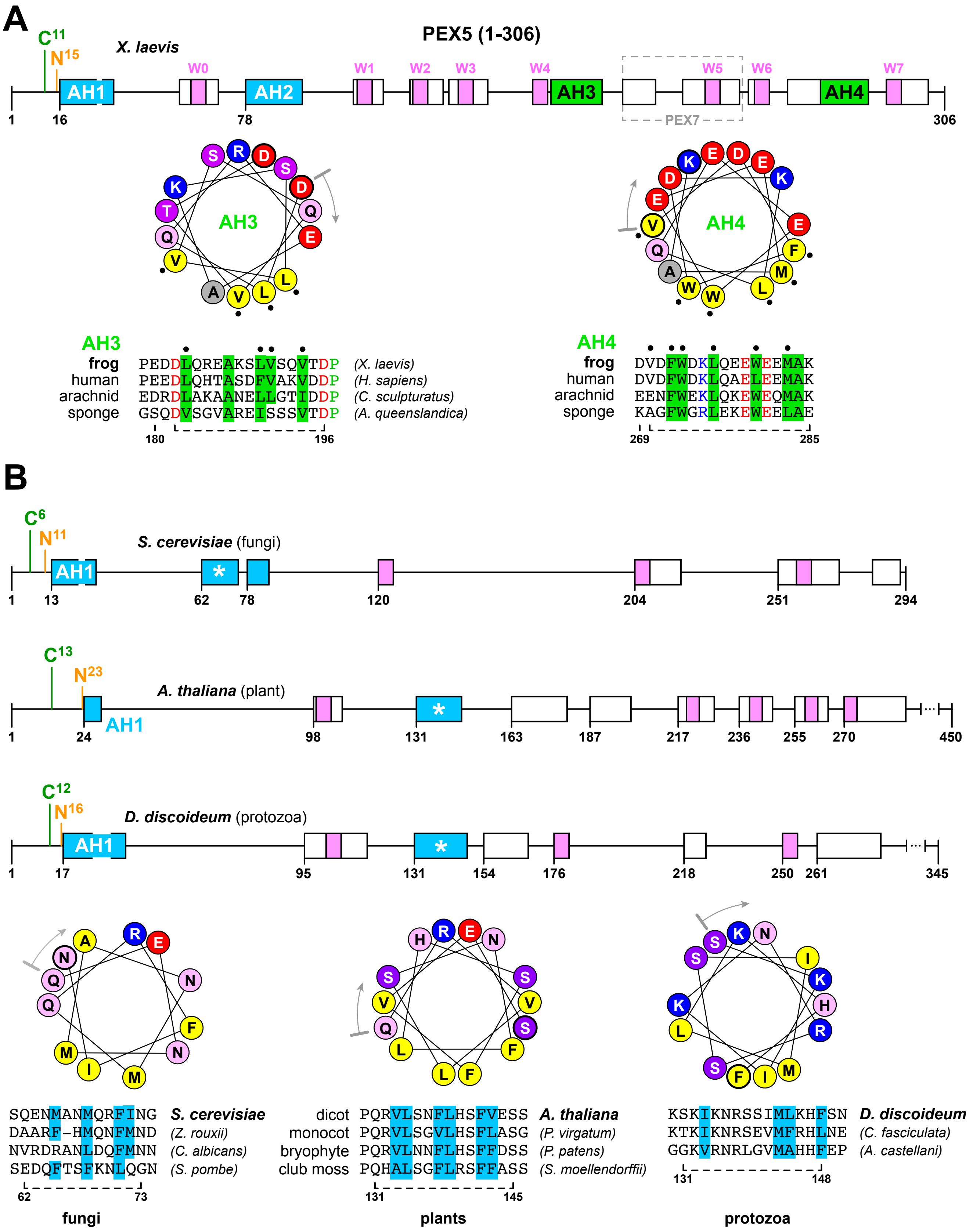

**Supplemental Figure S4. Conservation of amphipathic helices AH2-AH4 in the N-terminal unstructured region of PEX5 (related to Figure 3)**

(A) Diagram on top shows the locations of predicted α-helices (boxes) and key residues in the N-terminal unstructured region of *X. laevis* PEX5. Amphipathic helices are labeled AH1-AH4; WxxxF/Y pentapeptide motifs are labeled W0-W7. The PEX7-binding region is enclosed by a gray dashed line. Helical wheel diagrams illustrate the distribution of hydrophobic amino acids along AH3 and AH4 (gray arrows specify the N-terminus of each helix; black dots denote residues that were mutated to alanines). Underneath are sequence alignments of AH3 and AH4 in PEX5 homologs from the indicated organisms. Residue numbers refer to *X. laevis* PEX5; dashed line specifies the region shown in the helical wheels.

(B) As in (A), except that the N-terminal unstructured regions of PEX5 from the indicated organisms are shown. Amphipathic helices that might fulfill a similar role to AH2 in animals are labeled with an asterisk (*). Helical wheel diagrams illustrate the distribution of hydrophobic residues along each putative AH2 helix in PEX5 from the above organisms. The conservation of the hydrophobic residues is shown in the sequence alignments underneath. Dashed line indicates the regions used to construct the helical wheel diagrams; residue numbers refer to PEX5 from the organism in bold type.

### Supplemental Figure S5

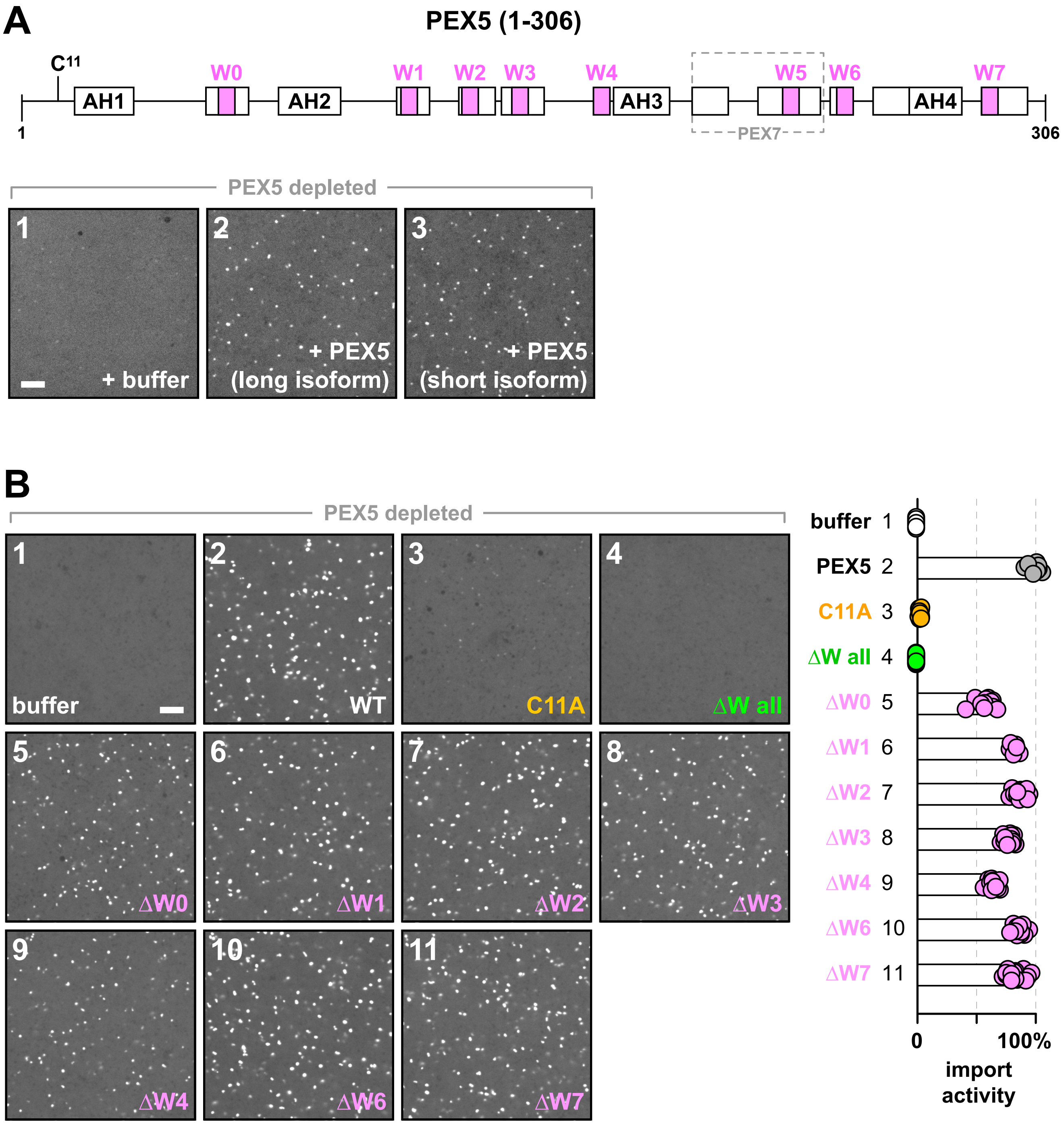

**Supplemental Figure S5. Import activity of PEX5 splice variants and mutants lacking individual WxxxF/Y motifs (related to Figure 3)**

(A) Diagram on top shows the locations of predicted α-helices (boxes) and key residues in the N-terminal unstructured region of *X. laevis* PEX5. Amphipathic helices are labeled AH1-AH4; WxxxF/Y pentapeptide motifs are labeled W0-W7. The PEX7-binding region is enclosed by a gray dashed line. Underneath, *Xenopus* egg extract was depleted of endogenous PEX5 using beads conjugated to the PEX5-binding domain from PEX14, then supplemented either with buffer, or with the long or short splice variants of PEX5. Import activity was assessed by incubating the reactions with GFP-SKL and imaging the formation of bright puncta on a spinning disk confocal microscope. The number of puncta in an imaged field was quantified relative to that in the reaction with wild type PEX5 ($n$ = 9 fields per reaction; bars specify the median).

(B) As in (A), except using the short PEX5 isoform (WT) or the short isoform containing the following mutations: Cys11 converted to alanine (C11A); all WxxxF/Y motifs mutated to AxxxA (ΔW all); or individual WxxxF/Y motifs mutated to AxxxA (ΔW0 through ΔW7). All scale bars equal 5 μm.

**Supplemental Figure S6**

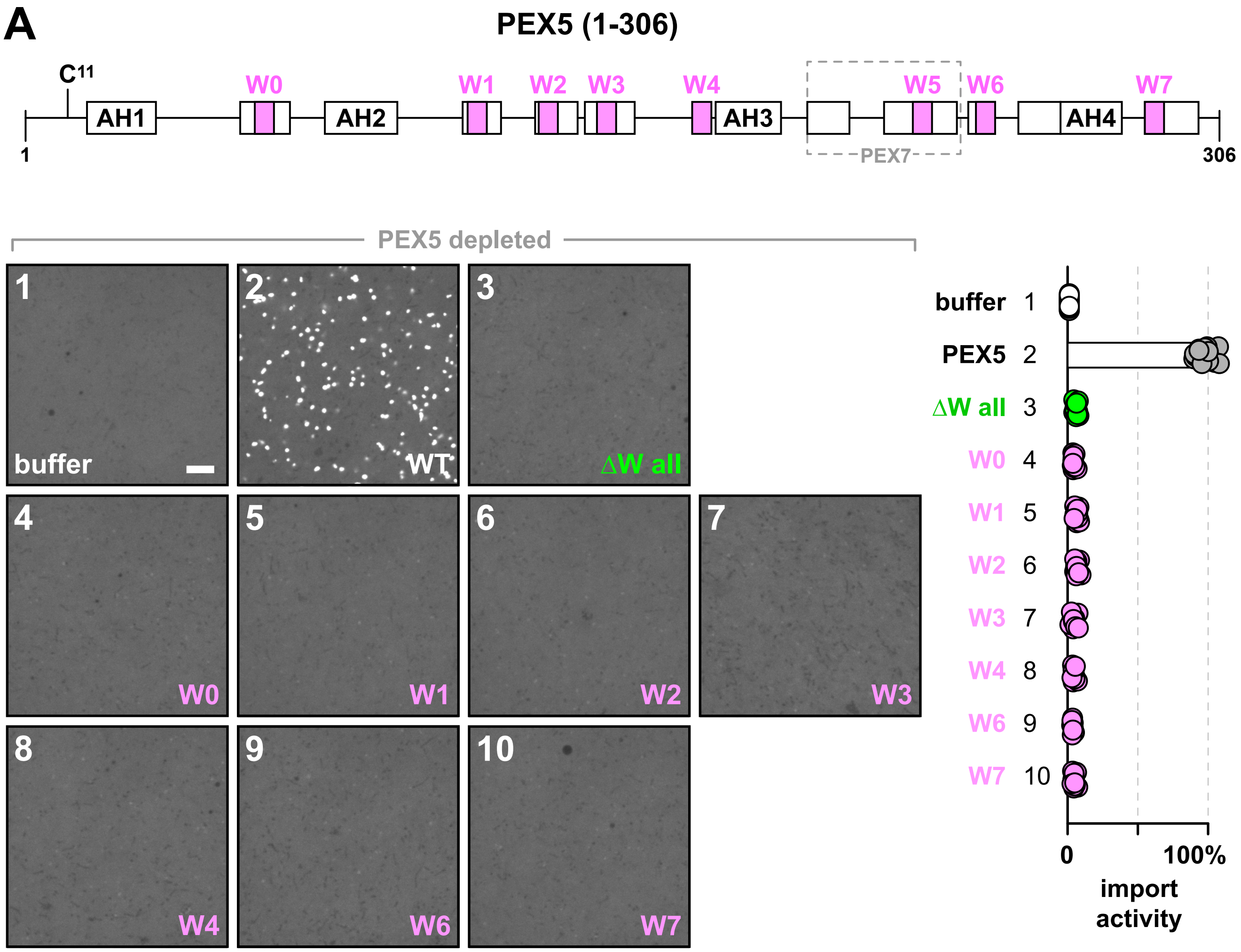


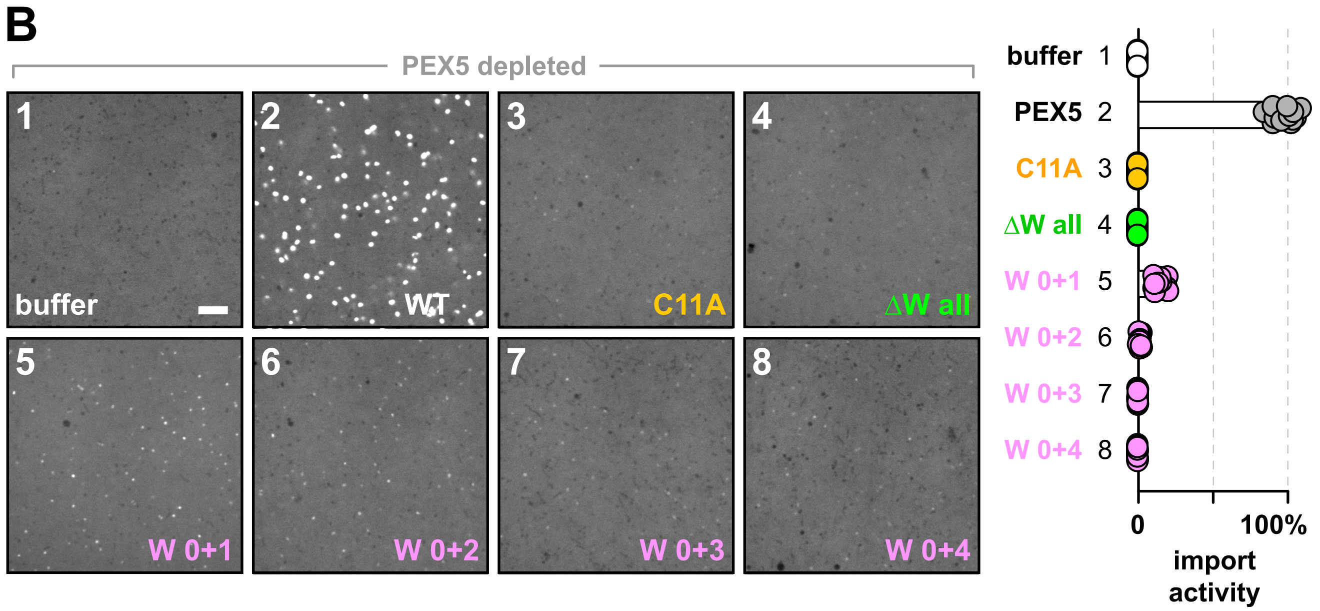

**Supplemental Figure S6. Import activity of PEX5 mutants containing different WxxxF/Y motifs (related to Figure 3)**

(A) Diagram on top shows the locations of predicted α-helices (boxes) and key residues in the N-terminal unstructured region of *X. laevis* PEX5. Amphipathic helices are labeled AH1-AH4; WxxxF/Y pentapeptide motifs are labeled W0-W7. The PEX7-binding region is enclosed by a gray dashed line. Underneath, *Xenopus* egg extract was depleted of endogenous PEX5 using beads conjugated to the PEX5-binding domain from PEX14, then supplemented either with buffer, wild type PEX5 (WT), a PEX5 mutant in which all WxxxF/Y motifs were converted to AxxxA (ΔW all), or PEX5 mutants containing only individual WxxxF/Y motifs. Import activity was assessed by incubating the reactions with GFP-SKL and imaging the formation of bright puncta on a spinning disk confocal microscope. The number of puncta in an imaged field was quantified relative to that in the reaction with wild type PEX5 ($n$ = 9 fields per reaction; bars specify the median).

(C) As in (B). C11A denotes a PEX5 mutant in which the conserved cysteine was mutated to alanine. Reactions in the bottom row were supplemented with PEX5 mutants containing only the indicated pairs of WxxxF/Y motifs. All scale bars equal 5 μm.

### Supplemental Figure S7

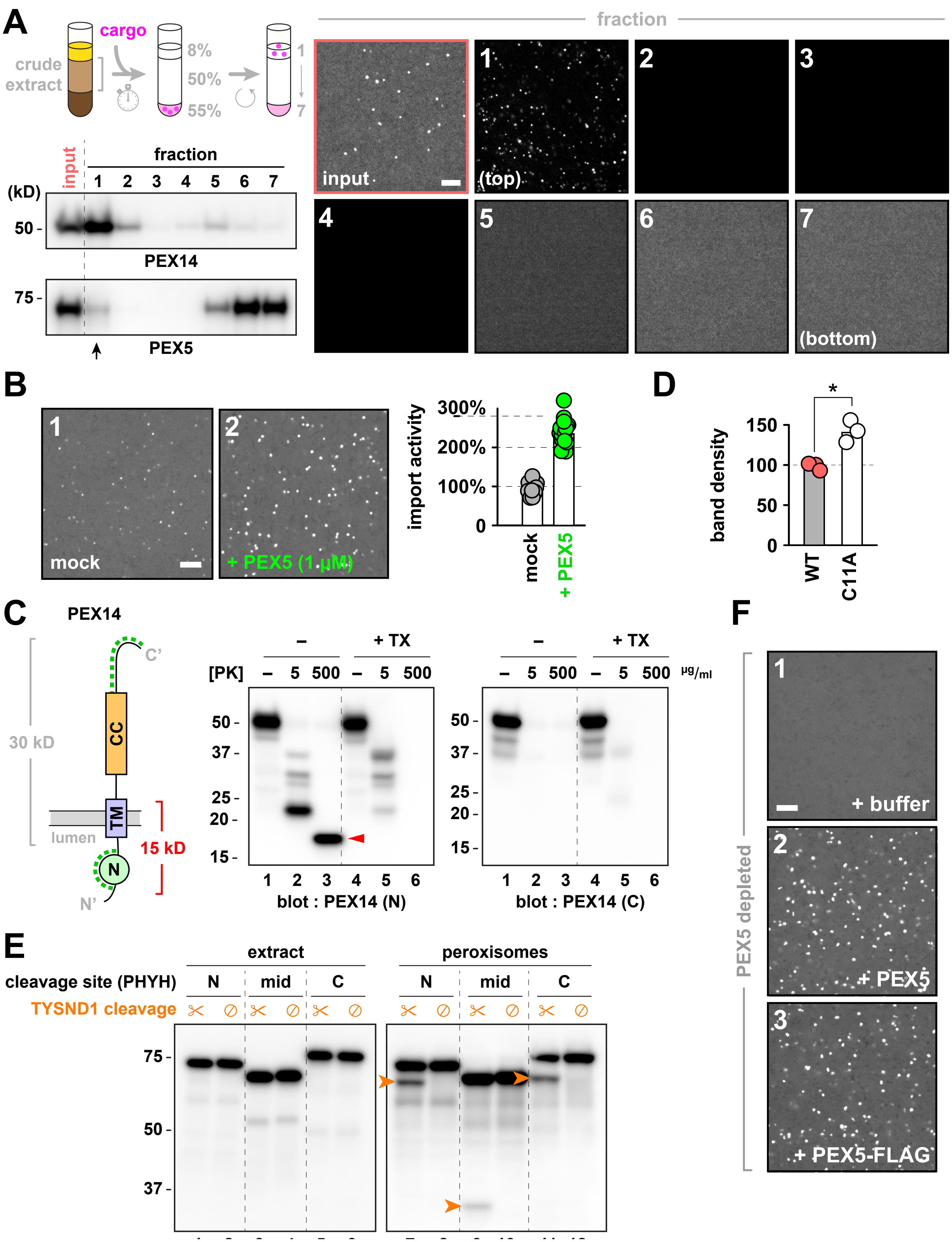

**Supplemental Figure S7. Validation of peroxisome isolation procedure, PEX14 topology analysis, and import activity of FLAG-tagged PEX5 (related to Figures 3, 4, and 5)**

(A) Scheme on top depicts the procedure for isolating peroxisomes from *Xenopus* egg extract. Peroxisome identification was facilitated by pre-incubating extract with mScarlet-SKL. Reactions were then diluted into a dense sucrose buffer and loaded on the bottom of a discontinuous sucrose gradient. Following centrifugation, peroxisomes accumulate on top of the gradient. On the right, fractions harvested from the top of the gradient were imaged by spinning disk confocal microscopy to confirm the separation of fluorescently-labeled peroxisomes (which appear as bright puncta) from cytoplasmic material containing unimported cargo (which appears diffuse). The starting material (input) is boxed in red. On the bottom left, the same fractions were immunoblotted for the indicated peroxins. Arrow indicates the peroxisome fraction.

(B) Extract was incubated with buffer (mock) or 1 μM recombinant wild type PEX5 in the presence of GFP-SKL, and formation of bright puncta was imaged on a spinning disk confocal microscope. The number of puncta in an imaged field was quantified relative to that in the mock reaction ($n$ = 9 fields per reaction; bars specify the median).

(C) Protease protection analysis of *X. laevis* PEX14. As shown in the scheme on the left, peroxisomes were isolated by flotation as in (A) and treated with different concentrations of proteinase K, with or without Triton X-100 (TX). Diagram in the middle depicts the orientation of PEX14 in the peroxisomal membrane, along with the predicted molecular weights of the indicated regions. Green dashed lines designate the segments used to raise polyclonal antibodies, which were then used on the right to immunoblot the protease-treated reactions. Red arrow indicates the expected protease-protected fragment.

(D) Quantitation of PEX5 density in lanes 3 and 6 of Fig. 4A, normalized to the density of PEX14 ($n$ = 3 independent experiments; *, $p \leq 0.01$ by Student's unpaired two-tailed $t$-test).

(E) As in Fig. 4F, except using wild type PEX5 (with the conserved cysteine intact) containing the TYSND1-cleavage site from PHYH at the indicated positions, as illustrated in Fig. 4C.

(F) Extract was depleted of endogenous PEX5 using beads conjugated to the PEX5-binding domain of PEX14, then supplemented either with buffer, wild type PEX5, or PEX5 fused at the C-terminus to a single FLAG tag. Import activity was evaluated as in (C), relative to the reaction with wild type PEX5. All scale bars equal 5 μm.

### Supplemental Table S1

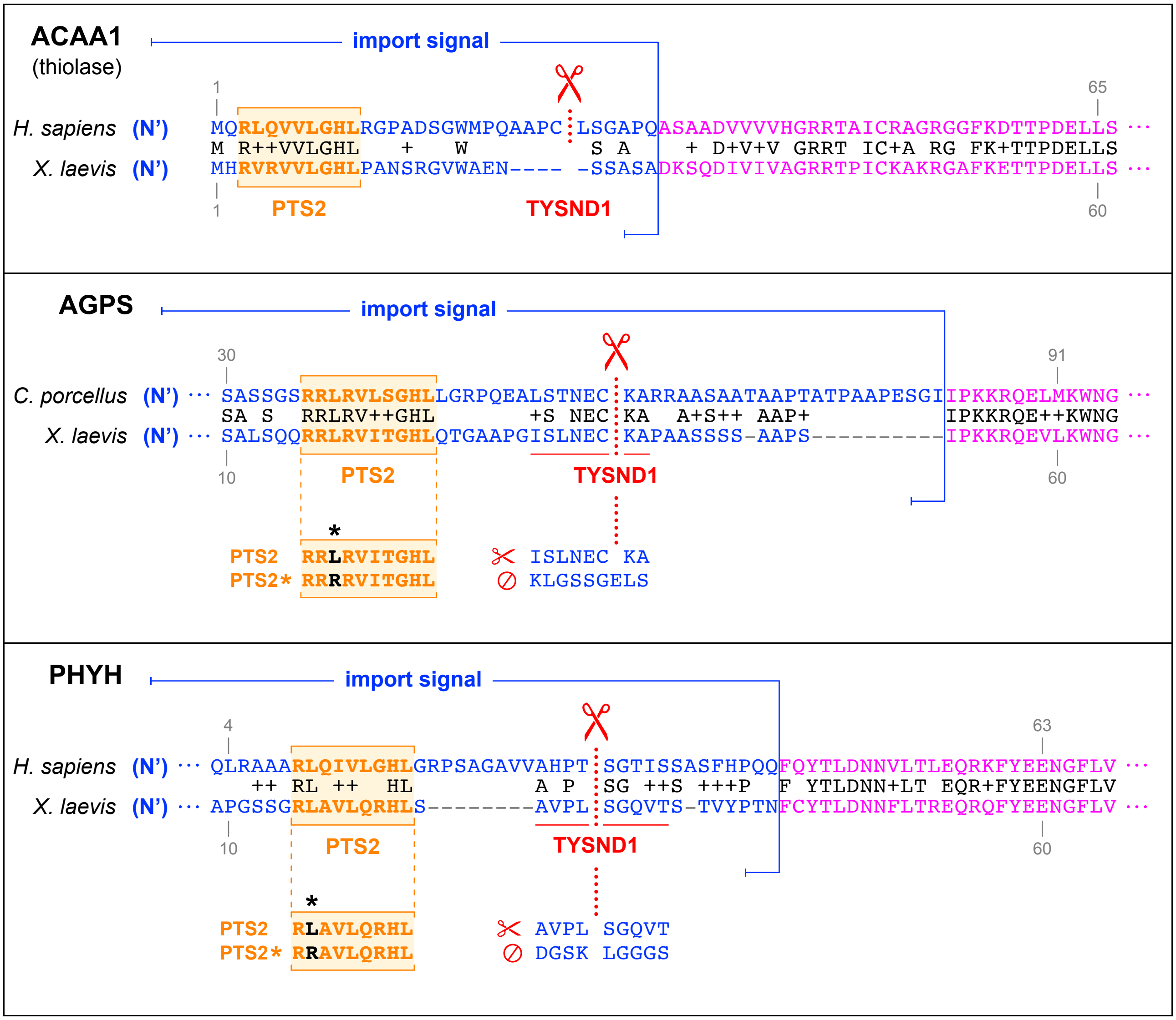


**Supplemental Table S1. Cleavable PTS2 signals used in this study (related to Figure 4)** The PTS2 import signals (blue) of the indicated peroxisomal matrix enzymes from mammals and *X. laevis* are shown (ACAA1, thiolase; AGPS, alkylglycerone phosphate synthase; PHYH, phytanyl-CoA hydroxylase). For each signal, the PTS2 motif is boxed and colored in orange, and the mutation used to inactivate the motif is indicated in black and marked with an asterisk (*). The TYSND1 cleavage site in mammals is indicated by scissors and a red dotted line. To inactivate the cleavage site in each *Xenopus* homolog, the residues underlined in red were mutated as shown underneath each signal. Note that the TYSND1 cleavage site does not appear to be conserved in *X. laevis* thiolase.
